# A parallelly distributed microscope and software system for scalable high-throughput multispectral 3D imaging

**DOI:** 10.1101/2025.05.31.657163

**Authors:** Hehai Jiang, Logan A Walker, Ye Li, Bin Duan, Xiaoman Niu, Jung-Chien Hsieh, Peng Yin, Mou-Chi Cheng, Hanquan Su, Kuanwei Sheng, Jessica P. Tang, Kenneth Athukorala, Stacey Greene, Ruby Pan, Arya Parlapalli, Meng Cui, Dawen Cai

## Abstract

Recent advances in high throughput optical microscopy have achieved whole-organism scale imaging at diffraction-limited resolutions. Current microscopes, however, require making compromises between achieving the optimal resolution, imaging depth, multispectral capability, and data throughput due to limitations in optical design and data stream handling. We have created a parallel-line scanning confocal microscope (plSCM) which provides 1.1 Gigavoxels/second high speed imaging while achieving an optical resolution of ∼180 ✕ 220 ✕ 650 nm with 2 millimeters of imaging depth in 3 simultaneous spectral channels. To handle such a massive imaging data stream, we have engineered a scalable network-distributed image acquisition/processing framework (SNDiF), which allows continuous capture, real-time processing, and cluster storage of petabyte-scale single image datasets in days. Together, our parallelly distributed microscope and software system presents a general solution for large-scale high-throughput and high-resolution multicolor imaging which can operate for days at a time.

## Introduction

Advances in large-scale and high-throughput imaging have revolutionized biological research, enabling unprecedented insights into complex biological systems. Whole-organ imaging allows for the comprehensive mapping of cellular and molecular architectures across entire tissues, providing critical context for developmental biology, disease progression, and organ-wide functional analysis^1,2^. Similarly, spatial-omics integrates molecular profiling with spatial localization, offering a high-dimensional view of cellular organization in diverse biological contexts^3,4^. These approaches have fueled breakthroughs in biology by linking cellular states with anatomical and functional structures. In neuroscience, connectomics, the systematic mapping of neural circuits, has become a powerful tool for understanding how brain structure underlies function^5–8^. Electron microscopy (EM) has been used to perform dense reconstruction of neuronal circuits in insect^9–11^ and mammalian^12,13^ brains. However, the high cost, prolonged imaging time, and limited imaging volume of EM constrain its applicability for large-scale tissue interrogation. In contrast, light microscopy (LM) is a more economical tool which is widely available in biomedical research labs. Yet, the reconstruction of neuron morphology at the whole-brain scale requires high spatial resolution to capture fine structures such as synapses, as well as large imaging volumes to be able to reconstruct entire projecting neurons^14–17^. Super resolution techniques, including expansion microscopy (ExM), have emerged as effective solutions for increasing achievable LM imaging resolution beyond the diffraction limit, but the inflated volumes of ExM samples necessitate faster imaging strategies to capture entire tissues in a reasonable timeframe. We have recently introduced spectral connectomics^18^ which allows neuron reconstruction using a combination of multiround ExM and Brainbow^19^ neuron labeling. However, this approach further demands highly efficient multispectral imaging.

Light-sheet microscopy (LSM) offers advantages for fast imaging but presents challenges in the sample mounting ergonomics and imaging millimeter/centimeter-sized samples with high NA objectives^20–22^. These limitations make LSM less flexible for imaging thick, expanded tissue slices. While recent developments in open-top LSM and oblique plane microscopy have improved sample mounting flexibility^23–25^, their axial resolutions are still above one micrometer in thick tissue samples, and multi-objective configuration requires precise alignments over long-term experiments. Lattice light sheet^26^ seeks to achieve high resolution by exploiting structured illumination shaping but is limited in imaging speed and sample size. Point scanning confocal microscopy remains a popular imaging method, valued for its high resolution, reliability, and adaptability across diverse experimental requirements, yet suffers from slow single-point scanning speed. Spinning disk confocal microscopy (SDCM) simultaneously scans hundreds of illumination spots on the sample, greatly improving the imaging speed. However, SDCM has limitations in simultaneous multi-color imaging due to spectral crosstalk and its associated shot-noise contaimination^27,28^, which restricts the applications of SDCM in high-throughput multi-spectral imaging.

Another challenge of high-throughput imaging systems is the process, transfer and storage of big data^29,30^. Modern sCMOS cameras enable high-speed biomedical imaging across very large fields of view with low noise, wide dynamic range, and high quantum efficiency across the visible spectrum with data rates of hundreds of megapixels to several gigapixels per second. Such a speed makes processing, storing, and visualizing of image data a major challenge. In one example^29^, an integrated imaging platform with a single sCMOS camera and a data management pipeline has been developed for quantitative data analysis with a throughput of 800 MB/s, matching the maximum data throughput of a single sCMOS camera. Handling data from multiple sCMOS cameras simultaneously, however, remains a major obstacle, especially for applications that require multispectral and high-resolution imaging at the whole organ/animal scale.

Here, we present a parallelly distributed microscope and software system to overcome the above-mentioned bottlenecks in scalable high-throughput multispectral 3D imaging. The system consists of two integrated components: (1) a high-speed multispectral parallel-line scanning confocal microscope (plSCM) and (2) a scalable network-distributed image acquisition framework (SNDiF). Leveraging our innovations in high aspect-ratio line illumination and chromatic aberration-free optical design, we simultaneously scan three spatially separated laser lines to excite fluorescent signals captured by three sCMOS cameras with minimized spectral crosstalk. This allows us to collect up to 1.1 Gigavoxels/sec at a spatial resolution of 180 ✕ 220 ✕ 650 nm^3^ in 2 mm imaging depth. At a voxel size of 105 ✕ 105 ✕ 250 nm, the plSCM is capable of capturing up to ∼220TB of data per day, corresponding to a volumetric throughput of ∼100 mm^3^ per day in three colors. Managing the high data throughput of up to 2.5 GB/s generated by plSCM motivated us to engineer SNDiF, a software package that enables continuous capture and cluster storage of PB-size single-image datasets. Unlike other microscopes, which rely on expensive SSDs for fast local storage, our SNDiF solution streams frames over a fiber optic network with real-time compression to quickly offload data onto a supercomputing environment or network filesystem. We demonstrate this parallelly distributed microscope and software system with multispectral super-resolution 3D imaging of different mouse and fruit fly brain tissues over millimeter-scale depths. This comprehensive system provides a versatile and scalable solution for high-throughput multi-spectral imaging.

## Results

### Parallel-line scanning confocal microscope (plSCM)

To minimize spectral crosstalk, plSCM simultaneously scans three spatially separated laser lines (488 nm, 560 nm, and 642 nm) (**Fig. 1a**, **Supp. Fig. 1**). Three 2304 ✕ 2304 sCMOS cameras (ORCA Fusion, Hamamatsu Photonics), each dedicated to a distinct spectral channel, are used to maximize data throughput. Combined with a high numerical aperture (NA) objective (Nikon CFI75 Apo25XC W, NA1.1), each camera supports a large field of view (FOV) of 242 ✕ 242 µm^2^ while maintaining high spatial resolution. To ensure diffraction-limited imaging, a high aspect-ratio (>1,500) excitation laser line is required to be precisely focused on the image plane across the entire FOV. In conventional line scanning designs, the excitation line is formed by a master cylindrical lens at the laser input and relayed to the back pupil plane of the objective^22,26^. However, to reach our desired high aspect ratio, a high-NA achromatic cylindrical lens would be needed, which is challenging to fabricate and susceptible to aberrations. We devised a simple solution by placing the master cylindrical lens C1 after the relay lens L4. With the fully expanded collimated laser beams, we could use a long focal-length cylindrical lens to achieve the desired aspect ratio without introducing aberration. The use of a long focal length lens also inherently reduces chromatic focal shift (**Supp. Fig. 2**). To completely eliminate the residual chromatic focal shift, we introduced an astigmatism control module (ACM) in the 560 nm and 642 nm laser paths, which comprises a pair of concave and convex lenses with a weak cylindrical lens in between (**Fig. 1b**). By adjusting distances S1 and S2, we can independently control the focusing or divergence in the tangential and sagittal planes, which allows all three laser lines to be precisely focused on the same image plane (**Supp. Fig. 3**). To minimize the wavefront distortion in the detection path, we positioned the dichroic beam splitters after the tube lens where the beam diameter is small and thus much less sensitive to the flatness error of the dichroic mirrors. Additionally, the smaller beam size on the dichroic mirror, the customized low-stress stainless steel dichroic mirror mounts, and the reduced distance between dichroics and the imaging sensors all contributed to minimizing long-term system mechanical drift. To avoid introducing astigmatism to the transmitted focused emission path, we used the dichroic mirrors at a small angle of incidence (21°), rather than the conventional 45°. Spectral modeling was employed to select the dichroic coatings such that their spectral cutoffs were at the desired wavelength at the designed incidence angle (**Supp. Fig. 4**). For volumetric recording, a fast piezo stage is used to acquire a 500 µm thick block, followed by a large-step movement with a 3-axis motorized stage to cover the entire 2 mm working distance of our objective (**Fig. 1e**). The single-objective architecture and the robust optomechanical design enabled high-speed, long-duration imaging of large and millimeter thick samples.

**Fig. 1.**
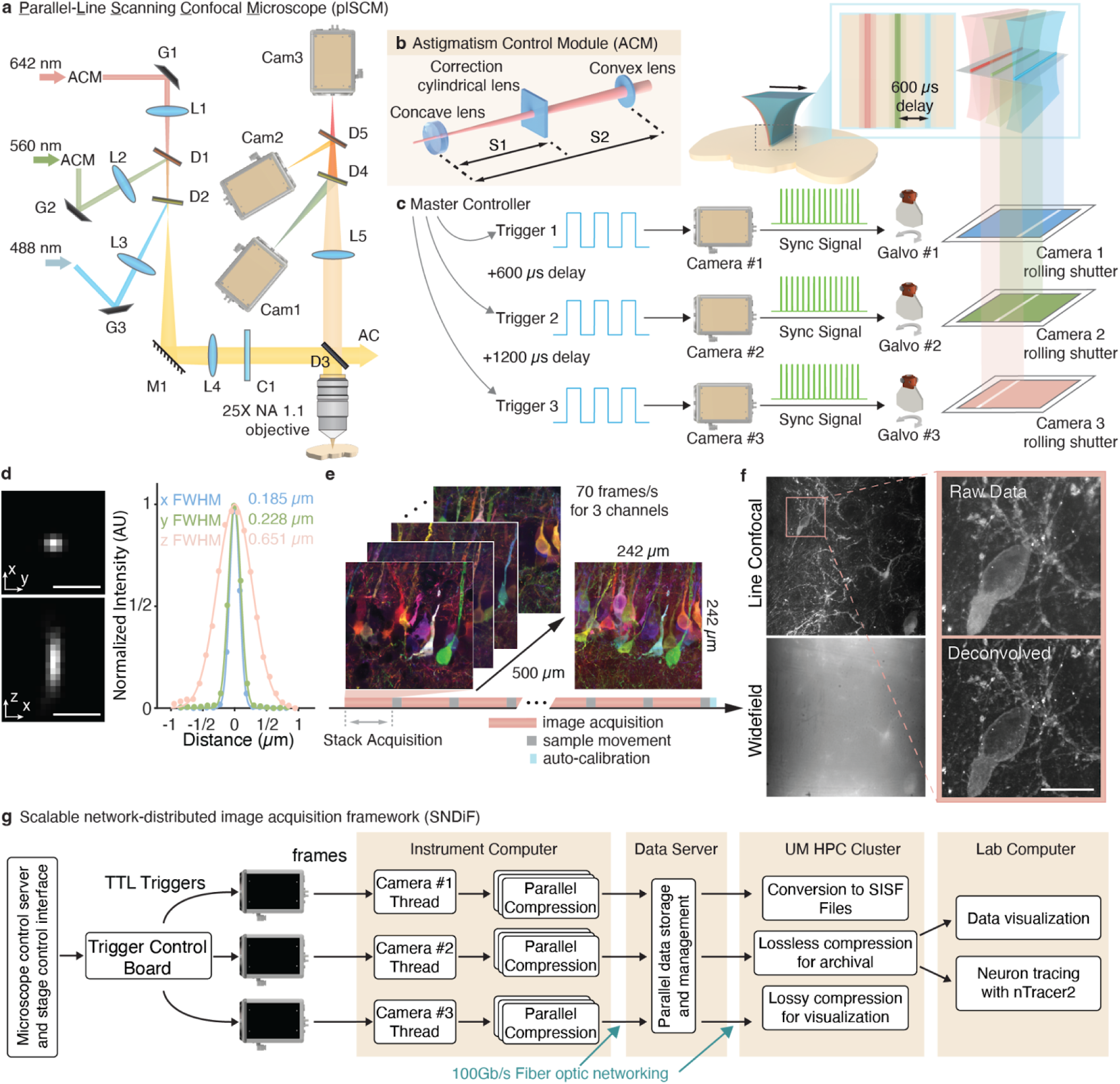
The parallelly distributed microscope and software system. a, Schematic of plSCM light path. G1-3, galvo scanners; L1-5, lenses; D1-5, dichroic mirrors; M1, mirror; C1, cylindrical lens. AC, auto-correction; Cam1-3, cameras. **b,** Outline of the astigmatism dual correction (ADC) module for maintaining high aspect ratio illumination. **c,** Overview of the imaging process. Each camera is triggered using an offset timing signal leading to physically-offset rolling shutters and line illuminations across the objective FOV. **d,** Representative point spread functions (PSF) from deconvoluted bead images with projections along *xy* and *xz* planes. Inset, FWHM measurements along each axis. **e,** Schematic of the timing sequence in a typical tile scanning experiment. **f,** Example GFP-labeled mouse neuron images with (upper left) and without (lower left) using the plSCM rolling shutter confocal mode. Zoomed-in views of the blue box are shown in the right panels. Both pre-deconvolution and post-deconvolution images are presented. **g**, Schematic of the scalable network-distributed image acquisition framework (SNDiF), wherein each camera’s data is collected and streamed in parallel, reducing the difficulty of handling the large amount of data.

The confocal detection is implemented through the rolling shutters of the sCMOS cameras. For each rolling scan, a custom-designed microscope controller^31^ generates three frame trigger signals with a 600-µs temporal delay, an equivalent of 13.6 µm in the image plane (**Fig. 1c** and **Supp. Fig. 5**) to each camera whose line clock synchronizes the scanning of its corresponding galvo (**Fig. 1c, inset**). In such a way, different laser lines are spatially separated, minimizing spectral crosstalk. For 3D optical sectioning, the width of the rolling shutter is set to 3.8 lines of pixels, creating an effective confocal slit of ∼24.8 µm, equivalent to ∼0.39 µm in the image plane. Precise synchronization between the rolling shutters and galvo scanning is crucial, as any mismatch between the rolling shutter and line focus can degrade contrast and resolution (**Supp. Fig. 6**). Previous solutions to this problem relied on visually adjusting the linear scanning voltage pattern to optimize imaging quality, which lacked quantity measurement and failed to account for the galvo scanner’s nonlinear response. To overcome this problem, we developed a high-precision calibration procedure (**Supp. Fig. 7**). To further mitigate potential drift during long-duration recording, we developed an automated tracking procedure using an auxiliary camera, which captures the scan pattern and compares it with the stored reference to quantify and correct the drift errors (**Supp. Fig. 8**).

To evaluate the spatial resolution, 200 nm fluorescent beads were mounted in a 1% agarose gel and imaged. After deconvolution with the bead profile, the result shows 183±6, 225±11, and 658±32 nm (μ±σ, n=137, **Fig. 1d** and **Supp. Fig. 9**) in three dimensions, respectively. To compare the performance with widefield imaging, we imaged a Thy1-GFP line M (GFP-M) mouse brain slice with miriEx^18^ expansion (**Fig. 1f**). The suppressed fluorescence background demonstrates the benefits of confocal detection on such thick expanded tissue samples.

### Scalable Data Handling

To handle the high data throughput produced by the plSCM cameras, we designed a scalable network-distributed image acquisition framework (SNDiF, **Fig. 1g**). SNDiF consists of three integrated components: first, a custom C++ application running on the imaging computer communicates to the cameras using the Hamamatsu application programming interface (API) to initiate the experiment and capture each image frame sequentially. Each frame is compressed using Zstd^32^ with byte-shuffling^33^ options and sent over the network to a custom server application which stores image stacks into a ZIP-based temporary format. Finally, a separate scripting interface running in Python is used to coordinate the tile scanning of the microscope. This combination of components allows streaming of extremely high imaging which we tested at 6 gigavoxels per second speed, which is way faster than all three cameras running at 80 FPS. Without needing to buffer data locally on the imaging computer, SNDiF takes advantage of the significant investments in centralized storage and compute capacity many institutions already have to ensure undisruptive high-speed imaging at the petabyte-scale.

### Large scale imaging of single cell morphology in the mouse brain

The high volumetric throughput combined with the single-objective architecture of the plSCM system make it ideal for large samples produced by expansion techniques, which can exceed 30X their initial volume^34^. To demonstrate this, we prepared a 0.5 mm thick GFP-M mouse brain slice using a modified miriEx^18^ protocol, achieving approximate 2.7X expansion (**Fig. 2a**). Native GFP fluorescence along with an antibody against GFAP (Rabbit anti-GFAP; Anti-Rabbit AF568), and DRAQ5 dye (ThermoFisher Scientific) were used to illustrate neurons, astrocytes and nuclei morphology, respectively. The resulting image was merged into a 108,000 ✕ 172,000 ✕ 2,000 (*xyz*) voxel image, consisting of 4,644 plSCM image tiles, with a 152 pixel overlap between tiles (**Fig. 2b; Supp. Vid. 1**). From the expansion factor, we estimate that each voxel had an effective ∼39 ✕ ∼39 ✕ ∼93 nm^3^ size. All image data was captured using SNDiF and converted into SISF files for visualization and annotation using the nTracer2 software platform^35^ and visualized using the nGauge Python package^36^.

**Fig. 2.**
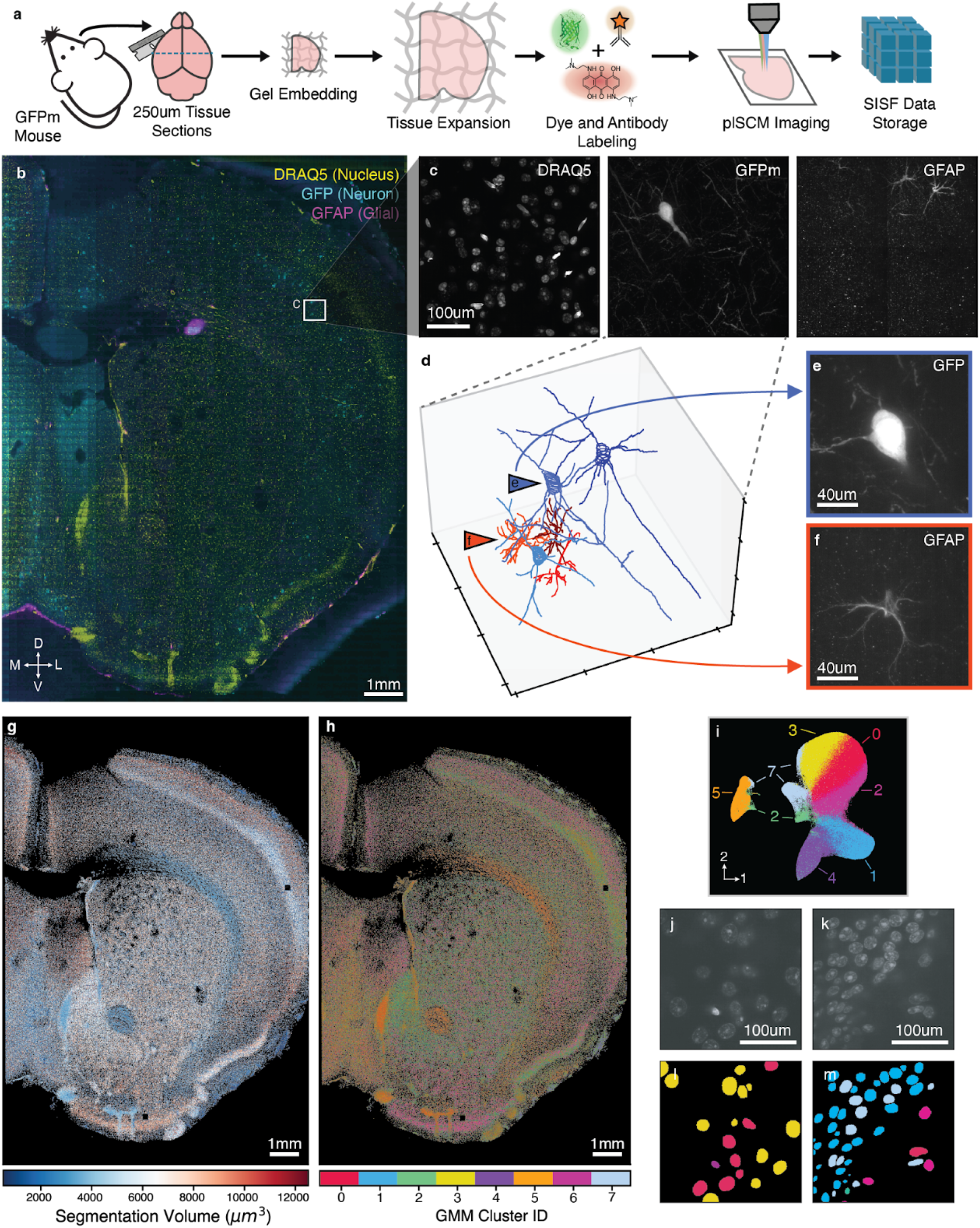
Super-resolution imaging at brain slice scale with plSCM. a, Outline of sample preparation protocol. **b**, Overview of one example Z-slice of a 2.7x expanded 250 µm thick hemi-section imaged in 54 ✕ 86 tiles (GFP neuron native fluorescence in cyan, DRAQ5 nuclei in yellow, GFAP astrocytes in magenta). **c**, Zoomed image of region indicated in **b**, displaying a maximum intensity projection of the three separate spectral channels. **d**, 3D rendering of neurons (blue tones, n=3) and astrocytes (red tones, n=3) which were reconstructed in the volume represented by **c**. **e**,**f** 50-frame maximum intensity projections (MIPs) of the GFP and GFAP channels, respectively, which correspond to two cells indicated in **d**. **g-h**, A scatterplot of each segmented DRAQ5-labeled nucleus in the sample, colored by the volume of the nucleus (**g**), or the GMM cluster identifier for each segmentation (**h**). **i**, A UMAP plot of each segmentation, colored by GMM cluster ID. **j-m**, Two zoomed views of DRAQ5 images (**j,k**) and the corresponding segmentations (**l,m**), colored by GMM cluster ID.

The resolution of the plSCM image allows us not only to identify individual neurons and astrocytes (**Fig. 2c; Supp. Vid. 2**), but also allowed us to reconstruct their refined morphology (**Fig. 2d-f**, n=3 GFP+ neurons, n=3 GFAP+ astrocytes). Using an nnUNET model^33^, we segment the 3D volumes of approximately 500,000 DRAQ5+ nuclei across the whole image. Using these segmentations, a variety of morphometrics were calculated for each nucleus (**Fig. 2g**, **Supp. Fig. 10**) and Uniform Manifold Approximation and Projection (UMAP)^37^ was used to perform dimensionality reduction on this data (**Supp. Fig. 11).** A Gaussian mixture model (GMM) was used to identify 8 morphometry clusters and their distribution on the brain section revealed interesting spatial patterns (**Fig. 2h-i, Supp. Fig. 12**), which indicates distinct brain cell types may be identified based on automated nucleus segmentation. Close inspection of the fluorescent images and their corresponding segmentations further confirm the distinct nuclei morphologies at the single nucleus level (**Fig. 2j-m**). Taken together, these analyses demonstrate the value of resolution and throughput of plSCM.

### Multispectral neuron imaging

Because the plSCM is optimized for simultaneous multispectral volumetric imaging, we hypothesized that it would be uniquely capable of imaging samples labeled by Brainbow^19,38^ and Bitbow^39^. To demonstrate this, we first performed imaging of expanded Bitbow *Drosophila Melanogaster* fruit fly brains (**Fig. 3**). TRH-Gal4^40^ flies were crossed with UAS-mBitbow2.0 flies^39^ to produce animals where serotonergic neurons are labeled with randomly mixed fluorescent proteins and their brains were processed with the miriEx^18^ expansion protocol (**Fig. 3a**). We expanded the gel to ∼2.4 fold, which results in a 13.8X volumetric dilution of the Bitbow labeling. Because plSCM cameras’ exposure times are very short, standard antibody staining results in images where neuron morphologies are difficult to fully reconstruct (data not shown). To create brighter samples, we utilized a novel Amplification by Cyclic Extension (ACE) protocol (detailed protocol unpublished, Yin lab) to enhance the antibody labeling intensity by more than 10-fold (**Fig. 3b**), resulting in substantially brighter labeling in the Bitbow mouse (**Supp. Fig. 14; Supp. Vid. 4**) and *Drosophila* brains (**Fig. 3c**, data not shown).

**Fig. 3.**
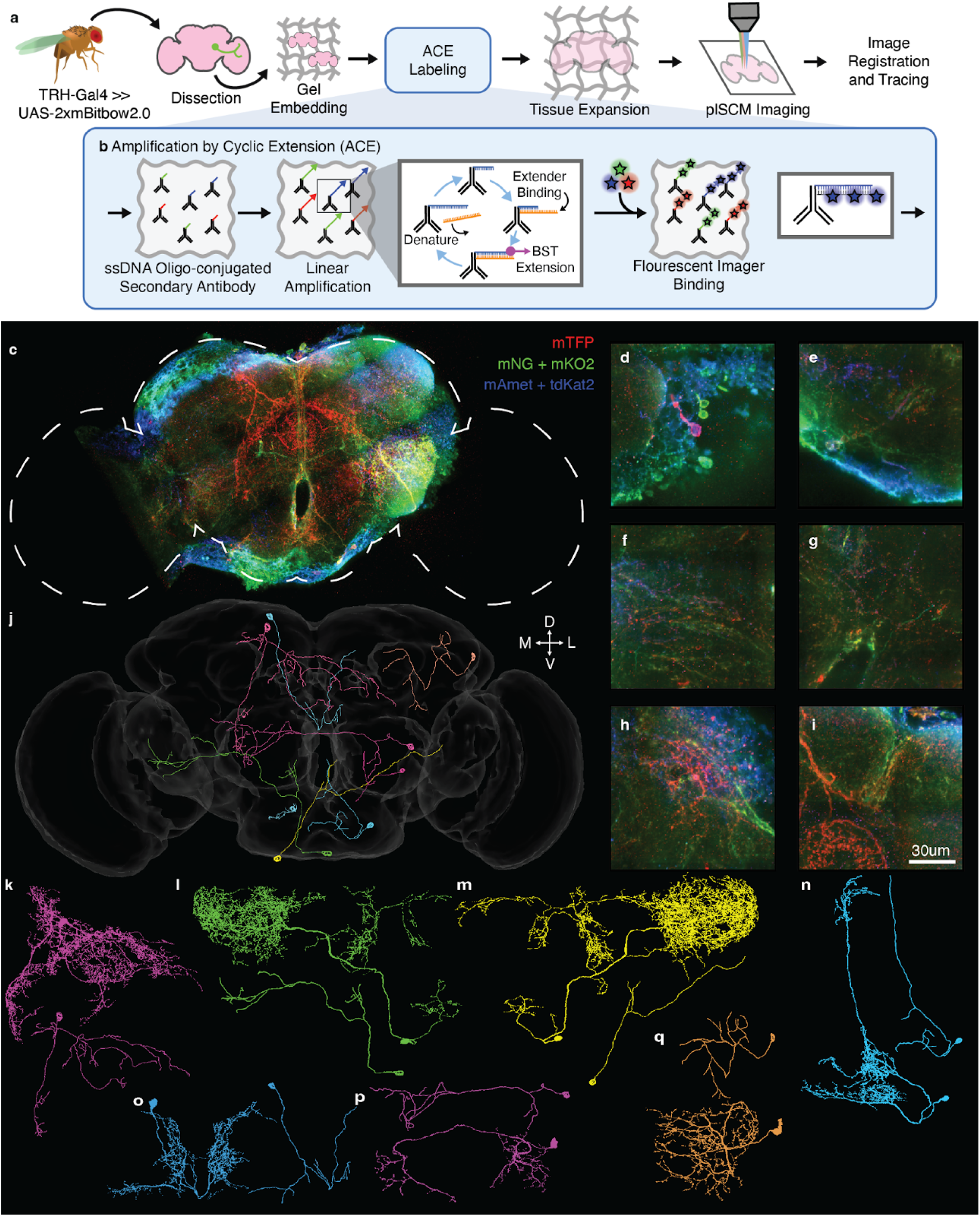
Multicolor plSCM enables high-speed Bitbow *Drosophila* imaging for neuron reconstruction and matching to EM reconstruction. **a**, Outline of sample preparation protocol. **b**, The amplification by cyclic extension (ACE) protocol is used to brightly label Bitbow antibodies using fluorescent oligos. **c**, Maximum intensity projection of serotonergic neurons labeled in a Bitbow *Drosophila* central brain. Outline represents the boundary of the JRC2018 fly brain template for anatomical reference. **d-i**, Zoomed-in images of different fly brain regions. **j**, Reconstructed morphologies of the Bitbow neurons traced using nTracer2 (n=9). **k-q**, Pair of matches identified between our reconstructions and the EM FlyWire reconstructions. The number at the bottom of each panel identifies the corresponding FlyWire Database ID s: 720575940610980932, 720575940637656974, 720575940618547641, 720575940622540899, 720575940621426300, 720575940630944843, and 720575940622474899.

We produced a 6,000 ✕ 6,000 ✕ 2,000 (*xyz*) voxel image, consisting of 3 ✕ 3 plSCM image tiles to cover most of the *Drosophila* central brain (**Fig. 3c; Supp. Vid. 3**). The resolution of the plSCM revealed great details of the serotonergic neurons (**Fig. 3d-i**, **Supp. Fig. 13**) and allowed us to perform 3D reconstruction (**Fig. 3j**) using nTracer2^35^. Next, we attempt to identify potential matches of our reconstructions to the FlyWire EM reconstructions^41^. To do so, we registered the overall brain contours of the FlyWire image volume (FAFB v783) to our plSCM image volume, followed by screening potential neuron reconstruction matches with similar soma locations and primary branching patterns using a custom pipeline (detailed in Methods). During our process of matching light microscopy (LM)-based traced neuron skeletons to the electron microscopy (EM)-based connectome library, we identified key morphological features that provided the highest value for extracting high-confidence matches. These critical features included: the first three orders of branches, long branches exceeding 100μm that cross neuropils, and soma location constrained within a 50μm radius. By prioritizing these features, we successfully identified 7 matched LM-EM neuron pairs from among the total 138,059 FlyWire reconstructions. **Fig. 3k-q** show 7 matching neuron pairs, which are also annotated as serotonergic or dopaminergic neurons in the FlyWire database.

### High speed total protein imaging

The LICONN protocol^17^ was recently reported, allowing researchers to achieve electron microscopy-like imaging contrast for connectomics studies in highly expanded brain samples. LICONN achieves this through a combination of multi-round sample expansion, total protein staining using N-hydroxysuccinimidyl (NHS)-esters, and machine learning segmentation of the resulting images. In the report, however, images were taken by a SDCM, which requires longer exposure times due to the low excitation efficiency. To investigate whether the plSCM could provide a solution for increasing the throughput of imaging highly expanded samples, we prepared a ∼10-fold expanded 50 µm mouse brain section using a similar protocol and performed single channel NHS-Cy3 imaging in the mouse cortex (**Fig. 4; Supp. Vids. 5-6**). As the effective voxel size falls in the 10nm range, we purposely slowed down the inter-tile stage moving speed to reduce motion artifacts. A 6 hr long, effective ∼45% duty cycle imaging session yielded a stitched 40,000 ✕ 40,000 ✕ 2,000 (*xyz*) voxel dataset that covers 4.2 mm ✕ 4.2 mm ✕ 500 µm imaging volume (**Fig. 4a**), which is a voxel rate of 191 megavoxels/sec (including stage movement and tile overlap). With 10X expansion, subcellular structures, including nucleoli, endoplasmic reticula, vesicles, lysosomes can be clearly resolved by our plSCM (**Fig. 4b-e**). Synapses can also be easily identified by our plSCM (**Fig. 4f-h**), similar to previously reported^17^, while gaining a 11.2-fold improvement of voxel throughput compared to using SDCM.

**Fig. 4.**
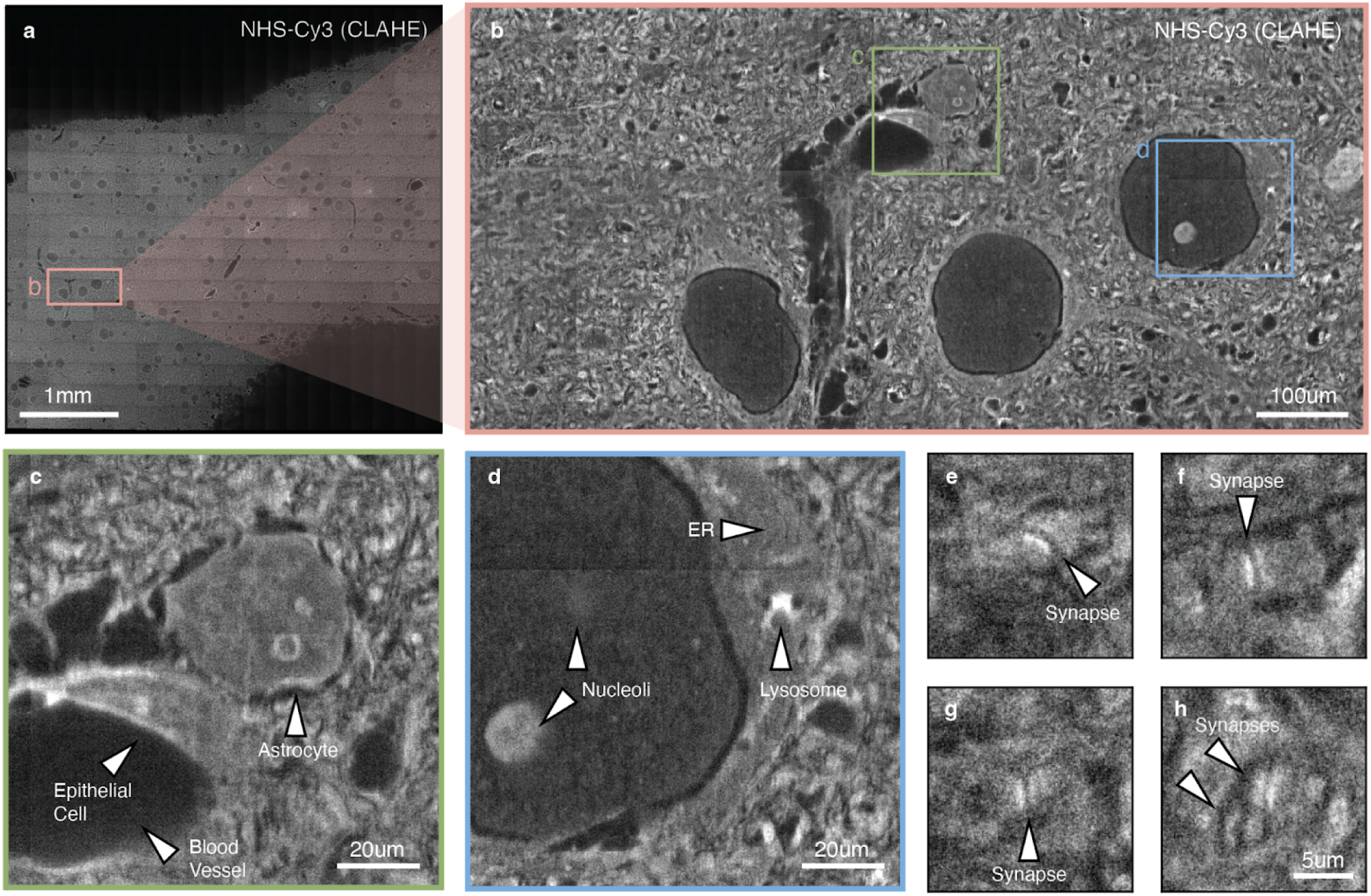
High-throughput 10x expansion imaging with the plSCM. **a**, Overview of N x M tile plSCM images covering a mm^3^ tissue volume. **b**, Detailed view of region indicated in **a**. **c-d**, Zoom-in views of regions indicated in **b**, annotating different putative brain cell types and cellular organelles. **e-h**, Zoom-in views of regions indicated in **b**, indicating putative synapses. All panels have been Guaissian filter smoothed and contrast adjusted using the Contrast-Limited Adaptive Histogram Equalization (CLAHE) method.

## Discussion

In this study, we have demonstrated a novel parallel-line scanning confocal microscope (plSCM) integrated with a scalable network-distributed image acquisition framework (SNDiF) that together overcome longstanding challenges in high-throughput, high-resolution 3D imaging. Our system achieves NA1.1 diffraction-limited line-confocal resolution over millimeter-scale imaging depths, providing a robust platform for large-scale neuroanatomical and tissue imaging studies. The optical design of the plSCM uniquely enables this through a novel high aspect-ratio line focus in conjunction with astigmatism dual correction (ADC), we effectively mitigate issues of chromatic aberration and spectral crosstalk that limit previous line confocal designs.

In parallel with these optical advancements, the development of SNDiF addresses one of the critical bottlenecks in high-throughput microscopy, the management of vast data streams. Our custom network-distributed acquisition framework seamlessly streams, compresses, and offloads data in real time to a supercomputing environment or network filesystem. This strategy circumvents the need for local high-speed storage solutions and enables continuous, long-term imaging sessions without data loss, paving the way for the routine capture of petabyte-scale datasets.

The versatility of our system is underscored by its successful application across diverse biological experiments. Imaging of expanded mouse brain tissues revealed intricate details of neuronal morphology, allowing for high-fidelity neuron reconstruction and the quantitative assessment of cellular features. Moreover, the capability to rapidly acquire multispectral images from Bitbow- and Brainbow-labeled samples demonstrates the plSCM system’s potential in complex multicolor labeling studies, essential for dissecting neural circuitry and cellular heterogeneity. Furthermore, the high-speed imaging of total protein stained LICONN samples underscores the system’s utility in accelerating light microscopy-based connectomics.

Despite these significant advancements, in this report we have only demonstrated a plSCM implementation which utilizes three laser lines. The concept, however, of using physically-separated line excitations can easily be logically extended to a near-unlimited number of laser lines, as long as matching dichroics, filters, and fluorescent dyes are available. Just using commodity computational hardware, our SNDiF software can handle multiple times of datastream than the current plSCM produces, which leaves an easy adaptation path for future microscope developments that utilize more and/or higher-speed cameras.

## Online Methods

### High-speed, Multispectral Parallel-Line Scanning Confocal Microscope

The plSCM was constructed on stainless steel breadboard using a list of parts available in **Supplemental Note 1**. In brief, a 488 nm laser (DL488-180-O, CrystaLaser), a 560 nm laser (VFL-P-300-560-OEM1-B1, MPB Communications Inc.), and a 642 nm laser (VFL-P-300-642-OEM1-B1, MPB Communications Inc.) were employed. An acousto-optic tunable filter (AOTF; AOTFnC-400.650-TN, AA Opto-Electronic) is used for power control. The laser beams traveled through galvo scanners (G1-3: 8320K, Cambridge Technology) and scan lenses (L1-3: f = 125 mm, two LA1301-A, Thorlabs) and were combined by two dichroic mirrors (D1: ZT561rdc-UF3, Chroma; D2: DI03-R473-t3, Semrock) before entering the tube lens (L4: f = 750 mm, AC508-750-A, Thorlabs). A cylindrical lens (CL1: f = 500 mm, LJ1144L2, Thorlabs) is inserted after the tube lens L4 to focus the laser beam into lines on the back pupil plane of a 1.1 NA imaging objective (CFI75 Apo 25XC W, Nikon). D3 (Di03-R405/488/561/635-t3, Semrock) serves as a multi-band master dichroic, which reflects the laser lines and transmits the fluorescent signals. The fluorescent signal collected by the objective travels through D3 and is focused by a tube lens (L5: f = 500 mm, AC508-500-A, Thorlabs) onto the cameras. The emission beam is spectrally separated into three paths by two dichroic mirrors (D4: DI03-R532-t3, Semrock; D5: DI03-R633-t3, Semrock) and simultaneously imaged by three 2304×2304 pixel sCMOS cameras (ORCA Fusion, Hamamatsu Photonics) equipped with bandpass filters (FF03-525/50; FF01-600/52; FF01-698/70, Semrock).

The sample stage is constructed from two 100 mm linear stages (UTS100PP, Newport Corporation) mounted transversely in X and Y, and a Z stage (9063-COM, Newport Corporation) with a motorized actuator (TRB25CC, Newport Corporation). These stages are controlled by two separate motor controllers (ESP302, Newport Corporation). An additional 500μm nanopositioning linear translation stage (P-625.1CD, Physik Instrumente, L.P.) is used to move the sample in Z rapidly during volumetric imaging.

### Microscope Automation

The imaging system was controlled through a custom Python 3.10 applet which interacts with the separate hardware components from a PC running Ubuntu Linux 20.04. Each galvo and laser power are controlled by our lab’s custom SimpleDAC microscope controller hardware^31^ which provides programmable signal generation, powered by three 30V power supplies (TP3005T, Tektronix).

Laser power, galvo synchronization, and movement of the stage are controlled by high frequency transistor-transistor logic (TTL) and serial signals between the respective hardware, controlled by Python scripts as detailed below. During imaging, each camera is triggered at a 70-80 Hz rate with a 600 µs offset between the start of each camera’s frame. As each camera exposes in rolling shutter mode, the VSYNC trigger output generates one pulse per pixel line and is used to synchronize the galvo motion using the SimpleDAC device. As the laser lines are aligned to the same physical position, this 600 µs initial trigger offset effectively scanned 600 µs / 4.6 µs/pixel (line rate) = 123 pixels on the camera, therefore, creating a (123 pixels x 6.5 µm/pixel size) / (62.5 total magnification) = 13.6 µm scanning offset at the sample between the illumination lines. To reduce photobleaching, at the end of each frame (flyback), each laser beam is shut off by the AOTF until the start of the next frame and the piezo stage is advanced by a transistor-transistor logic (TTL) signal. Each of these timing sequences is implemented as C code which is run on Raspberry Pi microcontrollers, with 5V logic shifting circuits used for compatibility with standard TTL signaling. Feedback from the piezo controller is used to detect if contact has been made with the objective to protect the microscope objective from damage and stops acquisition if the sample is out of position.

Each stage controller was connected through USB serial adapters (Prolific Technology, Inc.) and interacted with using the pySerial library [https://pypi.org/project/pyserial/]. The SimpleDAC and Raspberry Pi Pico used for shutter timing are connected over USB. A Xbox 360 game controller is used for stage control and positioning during experiment setup in lieu of a traditional joystick, with buttons mapped to trigger functions. During microscope operation, in a loop, the Python script checks for input from the game controller using the Python gamepad library [https://github.com/piborg/Gamepad] and translates the commands to the required stage commands. Additionally, a web API is implemented which allows stage operations to be actuated remotely. Using this, from a separate Python/Jupyter notebook, experiments can be defined using a simplified API which defines, e.g., the coordinates of a tile scan and relay them to the main microscope controller.

### Galvo Calibration Procedures

Because each of the galvo mirrors varies in their response to different driving voltages, calibration is required to determine the optimal control waveforms. This calibration is performed by first placing a fluorescent calibration sample (Chroma #92001) corresponding to the color of the laser being calibrated in the sample holder. A Python script moves each SimpleDAC to a list of defined voltages and a frame is taken at each location. From this information, a first order calibration is defined by linear extrapolation which provides a mapping between voltage and pixel location on the camera. This scan profile is uploaded to the SimpleDAC device and n=100 full-frame scans are taken. The residual of the intensity across this frame is used to further flatten the excitation profile by performing a polynomial fit and subtracting it from the first order calibration. This is referred to as the second-order calibration. Finally, this pattern is rotated by several pixels to confirm that it is a global maxima of excitation. This scan pattern is saved to the SimpleDAC for each channel and used for imaging. With the cylindrical lens rotated by a certain degree and a fluorescent pad positioned under the objective lens, we can analyze the galvo scanning pattern (**Supp. Fig. 6**). If the scanning is perfectly synchronized with the rolling shutter, the pattern observed on the camera is a straight line. Conversely, for scanning too fast or too slow, the line appears tilted. Should a mismatch between the scanning and rolling shutter speeds occur, lateral motion in the line will be observed.

### Stage Calibration Procedures

Because the X-Y stage is mounted on a separate platform from the rest of the instrument and may not be orthonormal, a 2D affine is performed between the targeted coordinates (location of image tiles) and the coordinates used to move the stage (physical locations). To calibrate this transform, a hematoxylin and eosin (H&E)-stained pathology slide is placed under the microscope. The stage is stepped 1 µm at a time in the X followed by Y directions, creating a library of several hundred images. Each of these images are registered together and the offset between frames is averaged to calculate a transform matrix which is then applied to future stage movements as a transform matrix.

### Optical Resolution Characterization

The microscope’s PSF was measured using 200 nm fluorescence beads across all three spectral channels (Thermo T7280). The beads are first embedded in agarose gel (1%) and then deposited into a petri dish. Three channel imaging was performed using a calibrated scan profile. The positions of the beads are then detected by finding the local maxima and cropping a volume around each bead for further analysis. The volumes are processed with 3 rounds of Richardson-Lucy (RL) deconvolution with the bead size. The PSFs are then fit with a one-dimensional Gaussian function along all three axes (x, y, and z) with a MATLAB script. This measurement was then confirmed using a similar procedure, but replacing the 200 nm beads with 100 nm beads (Thermo T7279), which provide a diffraction-limited PSF without the need for deconvolution, however at a much lower intensity.

### Automatic Drift Correction

During experiments which may continuously operate for days at a time, temperature and humidity drift in the microscope environment may cause small changes in the mechanical and optical calibration that reduces the image quality. To combat this, we mounted a calibration camera behind dichroic D3 (**Fig. 1a** and **Supp. Fig. 8**). This camera’s positioning allows it to capture excess light that is passed through the dichroic. Using this property, an automated on-the-fly calibration procedure is implemented which measures beam locations directly. Briefly, after performing the main galvo calibration described above, the galvo is swept while the laser is strobed every ∼10 pixels. During an experiment, the acquisition is paused and this scan pattern process is repeated for each laser to correct the drift of its corresponding galvo. Each peak in the strobe is identified and compared to the initial reference scan. Using the scan residual, changes in the galvo response can be measured and compensated for.

### SNDiF Implementation

The main SNDiF application is a C++ graphical user interface (GUI) program written in Microsoft Visual C++ 14 which captures individual frames from the Hamamatsu DCAM-API driver and performs Zstd^32^ compression. Each frame is uploaded to a server application written in the Rust programming language (compiled with rustc v1.65.0), which receives each frame in a scan and stores them in an individual ZIP container file. Because both the client and server application are multithreaded, multiple cameras can be handled in parallel and high network bandwidth can be achieved when combined with fiber-optic networking. A hard drive redundant array of independent disks (RAID) is used to capture the data from the SNDiF server.

### Image Preprocessing

The frames captured by SNDiF are converted one stack at a time using conversion Python scripts which perform a variety of preprocessing steps. First, frames are decompressed from both the ZIP and ZSTD container formats. The stacks of 2,000 frames are then combined into a single memory object representing the volumetric scan by sorting their timecodes. Next, frames are processed using BaSiCPy [https://pypi.org/project/BaSiCPy/], a python implementation of BaSiC^42^ for image flat field correction. To create the BaSiC model, a random sampling of n=200 frames from each channel of the experiment are selected which have mean intensity greater than average across the experiment and fit using BaSiCpy. We apply the resulting 2D model across all frames in parallel. Finally, each stack is saved as a downsampled image pyramid using the SISF image format^35^. Data processing is performed in parallel using SLURM or gnu-parallel^43^ for single-node jobs. The resulting archive is uploaded to an instance of the SISF-CDN^35^ for visualization and analysis.

We used a customized pipeline built upon Bigstitcher^44^ and the SISF file format, with a graphical user interface (GUI). This was used to stitch the Bitbow fly (**Figure 3**) and mouse cortex total protein staining (**Figure 4**) datasets. The documentation and code can be found at https://github.com/Cai-Lab-at-University-of-Michigan/stitching_gui/. For the Bitbow Fly dataset, channel registration was performed to align neurons labeled by different colors. Following the LICONN paper^17^, for the total protein staining dataset, we applied Contrast Limited Adaptive Histogram Equalization (CLAHE^45^) on the stitched image.

### DRAQ5 Stained Nuclei Segmentation

To perform segmentation of the nuclei image data (DRAQ5), we used a modified nnU-Net^46^ model. To increase computational throughput, we first downsampled the raw image data by a factor of 8x in the X and Y axes and 4× in the Z axis. We chose 12 representative tiles from the downsampled dataset and used Cellpose to obtain the initial segmentations, which were manually corrected for errors using nnInteractive^47^. The corrected segmentation was then used to train the customized nnU-Net. The morphometrics of the resulting whole-dataset segmentation were analyzed using customized scripts, which were then filtered for a segmentation being inside the volume outline, segmentation volumes (< 12,500 μm^3^), diameter (> 10 μm) and signal intensity (< 400 A.U.), to filter out outliers, which were verified to be segmentation errors by manual inspection. Each morphometric was combined into a single table, which was fit to a Gaussian Mixture Model (GMM) using scikit-learn^48^ version 1.5.1 and dimensionality reduction was performed with UMAP^37^ version 0.5.7.

### Fly Neuron Matching

The template “JRC2018” fly brain atlas^49^ outline from the Virtual Fly Brain^50^ (https://www.virtualflybrain.org/) and all EM neuron reconstructions from Flywire^41,51^ (https://flywire.ai/) were downloaded and manually transformed to align with the collected fly brain image inside of the neuroglancer interface [https://github.com/google/neuroglancer]. Flywire candidates were filtered for targets which were annotated as DA and SER neurons with a probability of >0.3 and then sorted by soma distance from our reconstructed TRH-Bitbow neurons. These candidates were visualized in batches of ten and manually screened by neuroanatomists to identify the highest-quality matches. For each target based on the primary branches.

### GFP-M Sample Preparation

4% PFA fixed coronal sections (500 µm) of an adult Thy1-GFP line M (JAX 007788) transgenic mouse brains were first permeabilized at 4°C with 10% CHAPS in MES-buffered saline (MBS, pH 6) for 24 hours. The samples were then incubated in 0.5 mM acrylic acid N-hydroxysuccinimide ester (AAx, Sigma A8060) in 1X MBS with 10% CHAPS overnight in the cold room. To complete the crosslinking reaction, the samples were washed with HEPES-buffered saline (HBS, pH 8.2) for 6 hours at room temperature (RT). Subsequently, the samples were incubated with a degassed monomer solution containing 8% acrylamide (Sigma-Aldrich A9099), 10.6% sodium acrylate, 0.01% bis-acrylamide (Bio-Rad, 1610142), 2% N,N-dimethylacrylamide (DMAA; Sigma 274135), and 0.5% VA-044 initiator (Fujifilm, 011-19365) in 1X PBS at 4°C overnight. It is crucial to ensure the solution is adequately degassed and kept cold to prevent premature polymerization.

The tissues were placed in a 500 µm iSpacer (SunJin Lab) on a slide, filled with the degassed monomer solution, and transferred to a humidified box for incubation at 37°C overnight to complete gel polymerization. Once gelled, the samples were removed from the spacers, excess embedding gel was trimmed, and washed with PBS for 1 hr at RT. The gelled samples were cleared using a modified miriEx protocol^18^. This was followed by washing in the clearing buffer for 1 hr and then in 1X PBS for another hour.

Cleared samples were protected from light and incubated in a blocking solution containing 5% donkey serum (JacksonImmunoResearch 017-000-121) and 0.5% Triton X-100 (Sigma-Aldrich 93443) at 4°C with gentle shaking overnight. The samples were then transferred into a staining buffer with polyclonal rabbit anti-glial fibrillary acidic protein (GFAP; Dako Z0334) antibody (1:500), 5% donkey serum, and 1% Triton X-100 in PBS. The tissues were gently shaken at 4°C for 3 days. After primary antibody staining (Dako Z033401-2), nonspecific binding was removed through three 20 min washes with 0.1% Triton X-100 in PBS at RT. A similar protocol was followed for three days of secondary antibody anti-rabbit Alexa Fluor 568 (1:500, Abcam ab175694) and DRAQ5 (1:1000, ThermoFisher 62251) staining. The stained samples were stored in PBS containing 0.02% sodium azide and kept at 4°C, protected from light, until ready for imaging.

### Bitbow Drosophila Sample Preparation

Flies were reared at 25°C on standard Caltech Medium food with a 12 hr/12 hr light/dark cycle. TRH-Gal4 Flies (Bloomington Drosophila Stock Center, BDSC 38388) were crossed with UAS-2xmBitbow2.0 flies^39^ (BDSC 99278) to produce animals where serotonin-producing neurons are labeled with random combinations of membrane-bound fluorescent proteins.

Three days after eclosion, adult brains were dissected in PBS at RT. Fixation was done with 4% PFA (EMS 15714) diluted in PBS with gentle nutation for 20 min, followed by three quick PBST (PBS + 1% Triton X-100) washes, then three PBS washes for 15 min. Afterwards, brains proceeded to one round of primary antibody staining, with sheep anti-EGFP, rat anti-mTFP, rabbit anti-mNeonGreen, mouse anti-mKusabiraOrange2, and guinea pig (GP) anti-TagRFP, before hydrogel embedding using miriEx^18^.

The miriEx gel-embedded brains went through another round of primary antibody staining with the same combination mentioned above and continued to ACE-specific secondary antibody staining, where DNA-conjugated antibodies p45-anti-sheep, p34-anti-rat, p40-anti-rabbit, p35-anti-mouse, and p53-anti-guinea-pig were used. Stained samples were fixed with 5mM BS(PEG)5 (Fisher 21581) at RT for 3h followed by quenching with 100mM glycine in 1X PBS at RT for 30 min. ACE amplification was done in the same fashion as described below, with a total of 500 amplification cycles. After amplification, the samples were incubated with 100nM DNA imager in 1X PBS for one overnight at RT, followed by six 20 min PBS washes at RT. Now, they were ready for mounting and imaging.

The hydrogel-embedded brains were slowly expanded to size, going through a series of gradual buffer exchanges from 1X PBS to 0.01x PBS. To reduce photobleaching, the samples were eventually placed in 0.01X PBS mixed with 20% (v/v) Vectashield PLUS (Vector lab H-1900), and mounted under a 22mm x 30mm No.1.5 coverslip (Corning 2980-223) coated with Poly-L-Lysine (Sigma P1524) spaced by a 1mm iSpacer (SunJin Lab IS012). The estimated expansion factor was ∼2.4× in the final mounting media.

### Bitbow Mouse Sample Preparation

A Flp and tTA double-dependent 6-color Bitbow1.0 mouse was crossed to CaMKII-tTA mouse (RRID:IMSR_JAX:007004). The double-positive offspring were injected with AAV9-hSyn1-Flp virus (10^12^ vg/ml x 800 nL) at 77 days old. After allowing five weeks for expression, the mouse was perfused with saline and 4% PFA and the brain was preserved in 30% sucrose (Sigma-Aldrich S0389). Two days later, the brain was sectioned into 100 μm slices.

To stain, slices were first incubated in 1% Triton X-100 (Sigma-Aldrich 93443) in 1X PBS for 4 hr at RT. Blocking was performed by incubating slices in 5% donkey serum (Jackson ImmunoResearch 017-000-121), 1% BSA (Sigma-Aldrich A3294), and 0.3% TritonX-100 in 1X PBS for 2 hr at RT. Primary antibodies were then added in blocking medium with gentle agitation at RT overnight. Antibodies used were hamster (Hm) anti-Myc (1:500; Absolute Antibody, Ab00100-22.0), rabbit (Rb) anti-mCherry (1:500, CancerTools 155260), mouse (Ms) anti-mKusabiraOrange2 (1:500; MBL Life Science, M168-3M-006), chicken (Ck) anti-mNeonGreen (1:250, CancerTools 155277), rat anti-mTFP1 (1:500, CancerTools 155264), guinea pig (GP) anti-TagRFP (1:500, CancerTools 155267). Each tissue was washed with 0.3% Triton X-100 in 1X PBS three times for 60 min. The buffer was then exchanged to 1X Modified Barth’s Saline (MBS) by incubating three times for 30 min.

To form the expansion gel, each section was incubated in 1.5 mM acrylic acid N-hydroxysuccinimide ester (AAx, Sigma A8060) and 0.1% Triton X-100 in 1X MBS at 4 °C overnight followed by washing with HEPES-buffered saline (HBS, pH 8.2) for 6 hours at room temperature (RT). Crosslinking quenching was performed by washing in 1X Tris-Buffered Saline (TBS; Bio-rad 1706435) three times for 1 hr at RT. After incubating in the monomer solution (same as GFP-M sample preparation above) overnight at 4 °C, slices were then placed on slides with a 0.15 mm iSpacer coverslip spacer (Sunjin Lab Co.) for gelling at 37 °C for 4 hr. Gelled samples were cleared by a modified miriEx^18^ protocol then washed in 0.3% Triton X-100 in 1X PBS at RT overnight. Each antibody was then restained at the same concentration as the initial staining above, followed by washing in 0.3% Triton X-100 in 1X PBS three times for 60 min. ACE staining was performed as described below prior to imaging.

### Amplification by Cyclic Extension (ACE) Protocol

Detailed ACE mechanism, barcode sequences, and antibody conjugation protocols were adapted from unpublished protocols developed in the Yin lab. In this study, the ACE protocol begins with a sample which has already been gel embedded and stained with primary antibody. To amplify a specific primary antibody, its secondary antibody is conjugated with an orthogonal ACE ssDNA barcode sequence The ACE secondary antibodies are diluted to 3ug/mL in 3% BSA, 0.2% Triton X-100, 0.02% dextran sulfate (Sigma-Aldrich D6001) and 500nM antibody DNA blocker, which are complementary ssDNA oligos to the barcode sequences. This solution is used to incubate the sample at 37 °C for 1 day. For fly and mouse Bitbow samples, secondary antibodies, anti-Rat-bc1, anti-Mouse-bc2, anti-Rabbit-bc3, anti-Guenea-Pig-bc4, anti-Chicken-bc5, anti-Hamster-bc6 were used accordingly. Tissues were then washed with 0.3% Triton X-100 in 1X PBS three times for 60 min. Postfixation was performed by incubation in BS(PEG)5 (Thermo-Fisher 21581), which was stored at -20 °C and diluted to 5mM in 1X PBS right before use. The sample was quenched by washing with 0.1M glycine (Sigma, G998-500G) in 1X PBS for 30 min. The sample is then washed with 30% formamide (Fisher Scientific BP228100) in 1X PBS overnight at RT, followed by rinsing in 1X PBS. ACE reaction was then performed for signal amplification (unpublished, Yin lab). After the ACE reaction, the sample is rinsed with 1X PBS followed by incubation in 60% formamide in 1X PBS overnight. Samples are again rinsed with 1X PBS and incubated with 1µM synthesized DNA imagers in 1X PBS for 2 hr, followed by washing in 1X PBS before imaging.

### 10x Expanded, Total Protein Stained Mouse Cortex Sample

Samples were prepared similarly to the previously published LICONN protocol to achieve well-preserved total protein staining and iterative 10x sample expansion^17^. An 8-week-old mouse was transcardially perfused with RT 1X phosphate-buffered saline (PBS) for 2 min, followed by a fixative solution containing 4% paraformaldehyde (PFA, pH 7.4, SC-281692) and 10% acrylamide (Sigma-Aldrich A9099) in 1X PBS. Brains were harvested and post-fixed for ≤12 hr at 4 °C with gentle agitation in the same fixative solution. Following postfixation, tissues were washed three times with cold (4 °C) 1X PBS for ∼1 min each with gentle agitation and stored in 1X PBS overnight at 4 °C. Coronal sections (50 µm) were obtained using a Vibratome (Leica VT 1200 S).

Individual sections were incubated in ice-cold 1X PBS supplemented with 100 mM glycine (Sigma, G998-500G) for 6-8 hr at 4 °C to quench reactive PFA groups, followed by three 1-min washes in 4 °C 1X PBS and storage in 1X PBS with 0.015% sodium azide (ThermoFisher Scientific 71448-16). Protein anchoring was performed by first washing each section twice for 15 minutes in RT PBS with gentle agitation. Each section was then washed twice for 15 minutes in 100mM sodium bicarbonate in ddH_2_0 (pH 8.0) at RT. Sections were then incubated in a protein anchoring solution (1% w/v glycidyl methacrylate [GMA; Sigma-Aldrich 779342] and 1% w/v triethylene glycol dimethacrylate [TGE; Biosynth FG173064] in 100 mM sodium bicarbonate, pH 8.0) for 3 hr at 37 °C with gentle agitation in a fume hood. Finally, sections were washed in 1X PBS for 1hr with gentle agitation at RT.

The first expansion hydrogel was prepared by mixing 10% (w/v) acrylamide, 12.5% (w/v) sodium acrylate (SA, Sigma-Aldrich 408220), and 0.075% (w/v) bis-acrylamide (BIS; Bio-Rad, 1610142) in ddH_2_O and supplemented with 0.5% (w/v) VA-044 initiator (Fujifilm, 011-19365) on ice. Degassed monomer solution was used to cast sections with a 0.15 mm spacer (Sunjin Lab Co.) between two glass slides, incubating at 4 °C overnight in a humidified, dark chamber, followed by a 2 hr incubation at 37 °C in a light-sealed container. Post-gelation, hydrogels were trimmed to desired regions of interest and washed three times in 1X PBS. Gels were transferred to a denaturation buffer (200 mM sodium dodecyl sulfate [SDS, Bio-Rad 1610302], 200 mM NaCl [ThermoFisher Scientific S271-1], 50 mM Tris-HCl [invitrogen, 15567-027], adjusted to pH 9.0 with NaOH), and autoclaved at 120 °C for 30 min. Samples were washed with PBST (0.1% Triton X-100 in PBS, Sigma-Aldrich 93443) three times and expanded in ddH_2_O. The water is changed every ∼20 min until no further expansion is observed. Each gel is then incubated in 1% VA-044 in dH_2_0 to neutralize unreacted BIS in 37 °C for 1 hr, followed by washing in ddH_2_O, first in 37 °C for 30 min, followed by RT for 30 min. The second expansion was performed similar to the first expansion except using 10% (w/v) AA, 19% (w/v) SA, and 0.025% (w/v) BIS in a chamber made with 0.5 mm spacer.

Total protein restaining is performed by incubating each sample in a 40 μM Cy3 NHS-ester solution in 1X PBS overnight at 4 °C with gentle agitation. The slide is washed with 1X PBS, three times for 5 min, followed by expansion in ddH_2_O at RT for 30 min. For imaging, each gel was mounted on a slide and 1mm iSpacer coverslip spacers were used to mount a coverslip over the tissue without pressing on the gel, which would cause deformation.

## Supplementary Figures

**Fig. S1.**
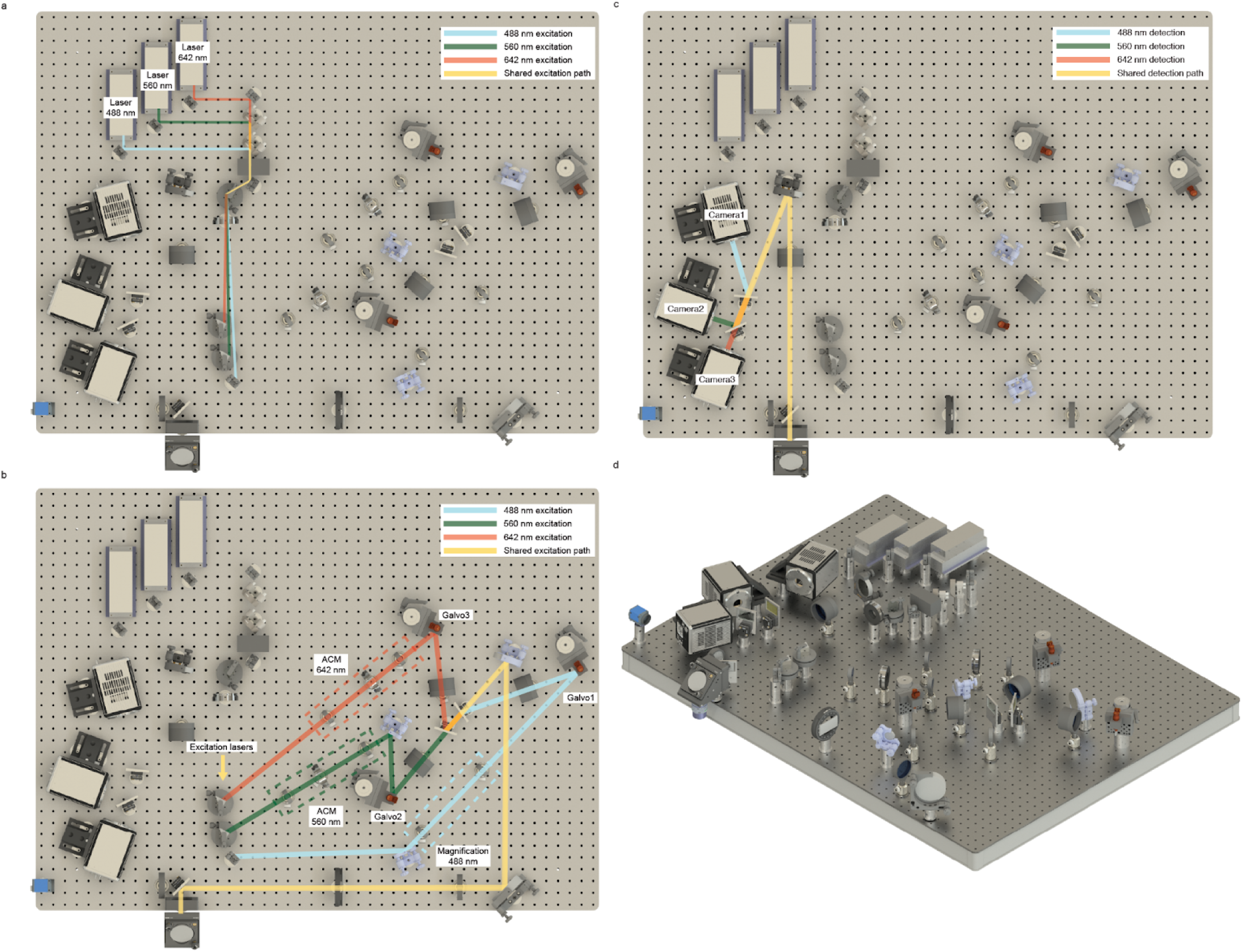
CAD and mechanical design of the plSCM. All components are mounted on a 36 ✕ 48 inch breadboard. **a-b,** Top-view of the excitation paths. Laser beams at 488 nm, 560 nm, and 642 nm are modulated by an acousto-optic tunable filter (AOTF), followed by a prism to separate three wavelengths into each ADC and galvo scanners. Three laser beams are subsequently combined by dichroics and directed into the objective lens. **c,** Top-view of the detection path. The fluorescent signals collected by the objective lens are spectrally separated into three independent cameras, enabling simultaneous multi-color imaging. **d,** Overview rendering at 45° view.

**Fig. S2.**
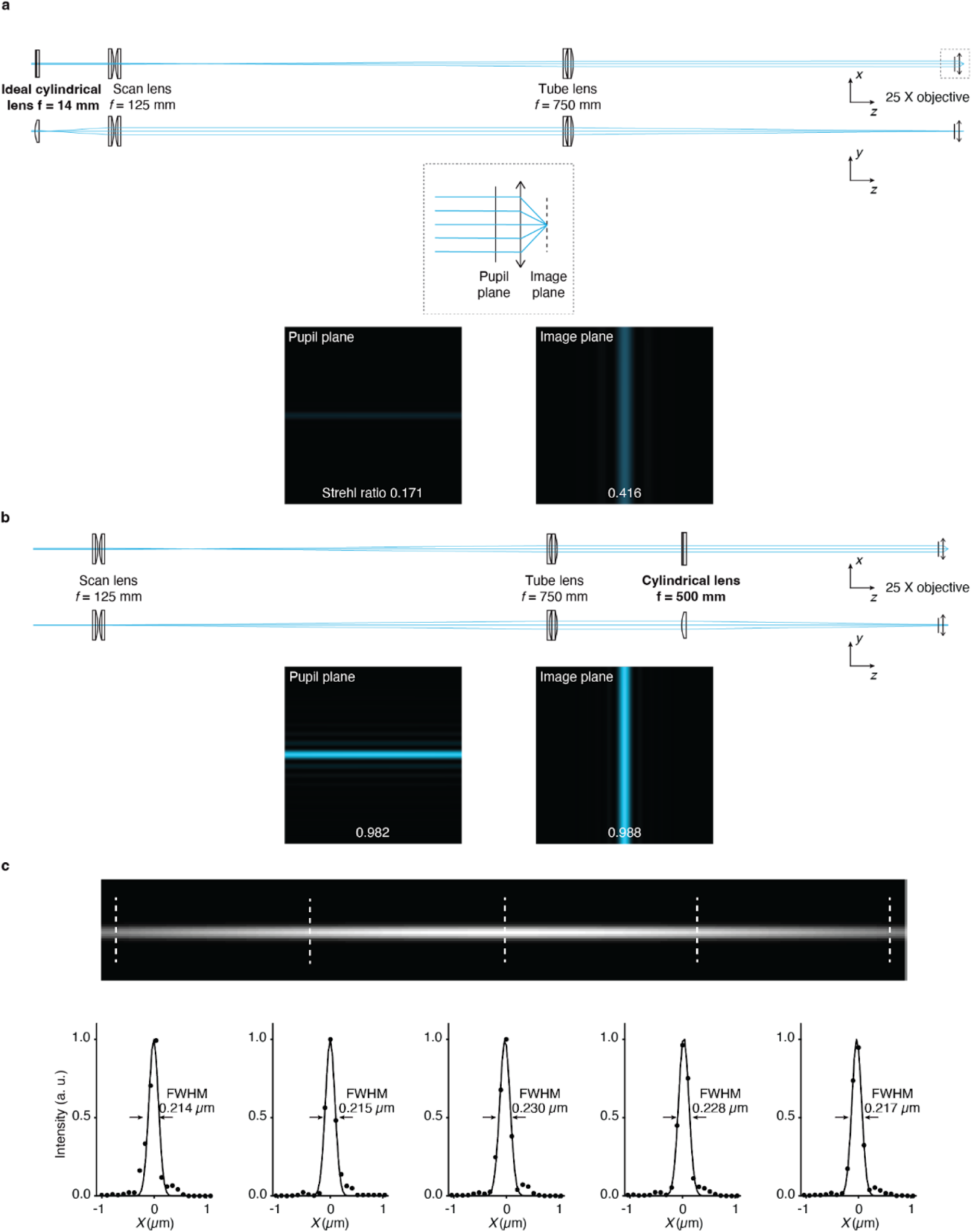
plSCM excitation path optical design. a, Conventional optical design directly elongates the Guassian laser beam to a scanning line by a cylindrical lens and then projects it through the relay pair onto the pupil plane. Dashed box, zoom-in of the region around the objective lens, showing the pupil plane and the image plane. Lower panels, the Huygens PSF at the pupil plane and the image plane indicates a low quality scanning line formed by the conventional design. **b,** plSCM design first expands the Guassian laser beam by the relay pair and then elongates it by a cylindrical lens and projects onto the pupil plane. Lower panels, Huygens PSF at the pupil plane and the image plane indicate a high quality scanning line formed by the plSCM design. **c,** Measured plSCM excitation PSF at 5 different positions along the line illuminations. The upper panel renders a simulated line profile to better indicate the PSF positions.

**Fig. S3.**
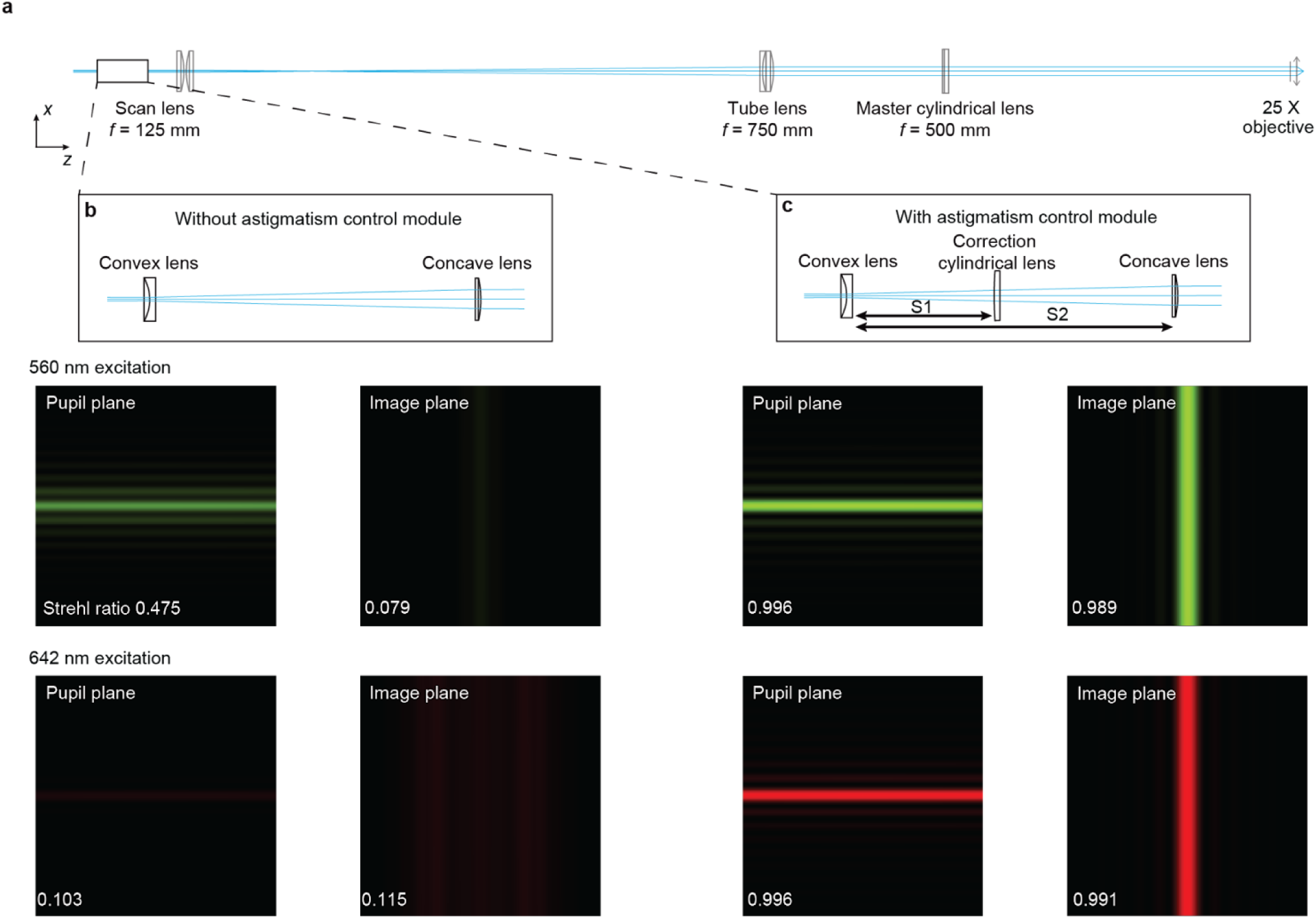
Astigmatism dual correction (ADC) to correct chromatic aberration introduced by the cylindrical lens. a, Optical layout of plSCM excitation line design. **b,** Without ADC, the convex/concave lens pair is used solely for beam magnification. There is significant chromatic aberration introduced by the master cylindrical lens. Upper panels, Huygens PSF at the pupil plane and the image plane of 560 nm. Lower panels, Huygens PSF at the pupil plane and the image plane of 642 nm. **c,** ADC design introduces an additional correction cylindrical lens to correct the chromatic aberration for each laser light path. Upper panels, Huygens PSF at the pupil plane and the image plane of 560 nm. Lower panels, Huygens PSF at the pupil plane and the image plane of 642 nm. The near perfect Strehl ratios indicate chromatic aberrations have been independently corrected for each laser.

**Fig. S4.**
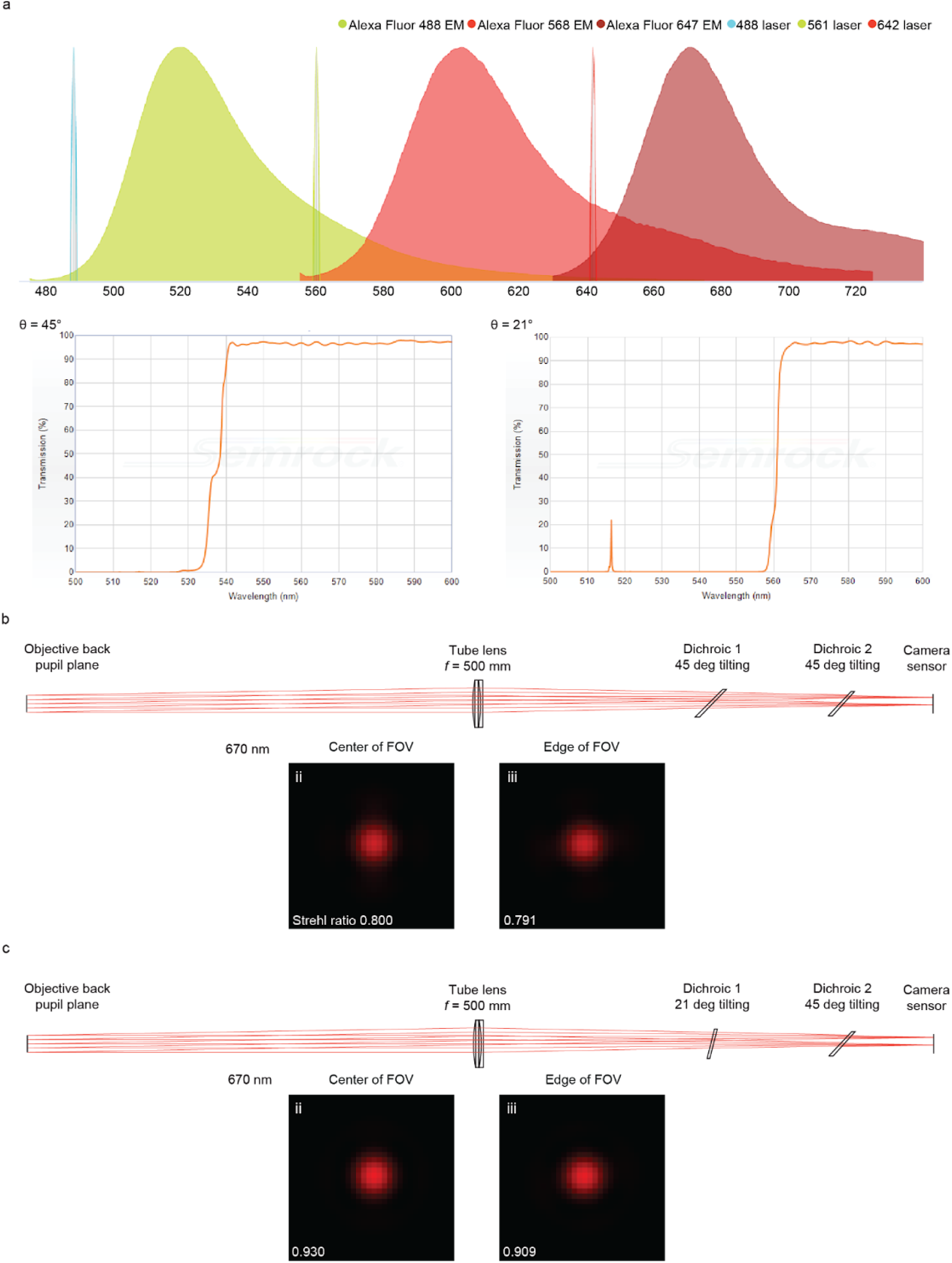
Tilting angle of the dichroic mirror affects its transmission spectrum and the focus quality. a, Upper panel, fluorophore spectrum and lasers used in the experiment (source: https://www.fpbase.org/). Lower left or right panel, transmission spectrum of dichroic mirror DI03-R532-t3 positioned at a 45° or 21° incident angle, respectively (source: https://www.idex-hs.com/store/product-detail). **b**, When Alexa Fluor 647 emission passes both dichroic mirrors (DMs) positioned at 45° incident angle, its astigmatism significantly diminishes the focus quality on Camera 3 sensor. **c**, When the first DM, DI03-R532-t3, is positioned at 21° incident angle, Alexa Fluor 647’s focus quality on Camera 3 sensor is greatly improved.

**Fig. S5.**
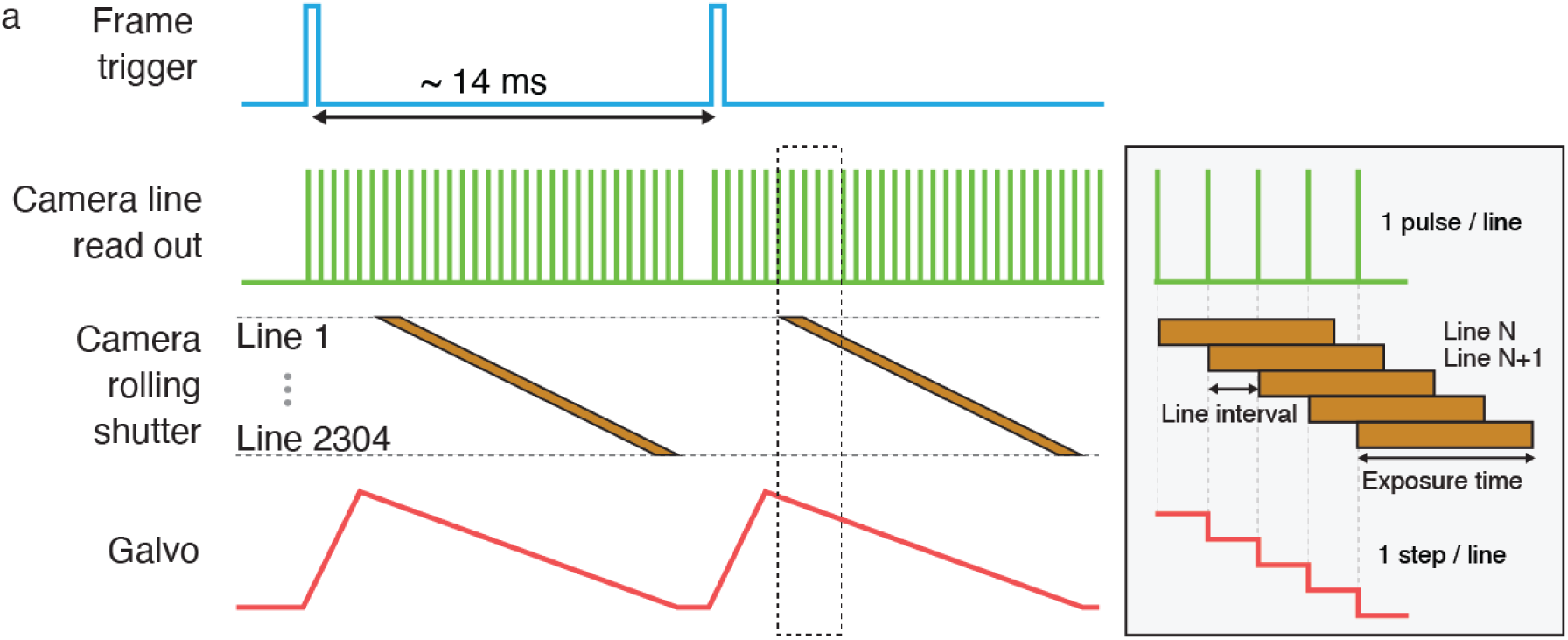
plSCM electromechanical timing. a, plSCM trigger timing. The frame trigger starts the camera’s rolling shutter, while simultaneously outputting the camera’s line readout signal, which serves as the clock for synchronizing other instruments. With the clock signal, the galvo scanners start scanning and the lasers operate at a constant intensity and are switched off between frames by the AOTF. The *z*-axis piezo stage advances by one step between each frame for 3D volumetric imaging.

**Fig. S6.**
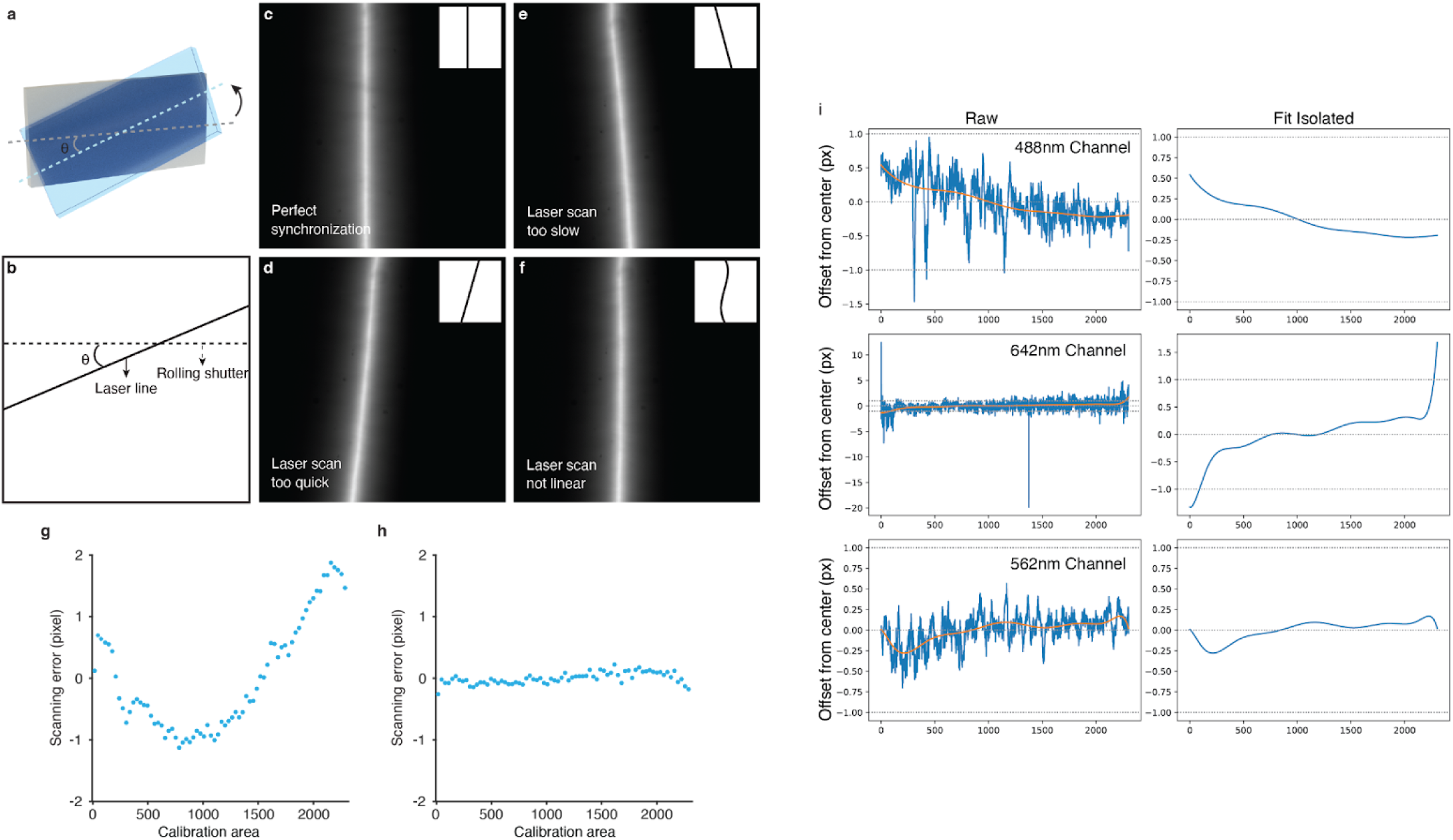
Galvo and camera synchronization impact on the imaging quality. a, Maximum-intensity projection (MIP) of 0.2 µm fluorescent beads in 1% agarose gel with synchronized galvo scanning and camera rolling shutter. Inset shows the Fourier transform of the image. **b,** MIP of 0.2 µm fluorescent beads with a 6 px offset between galvo scanning and camera rolling shutter. Inset shows the Fourier transform of the image. Scale bars, 100 µm. **c,** Magnified view of white dashed rectangle region with 0/2/4/6 px offset.

**Fig. S7.**
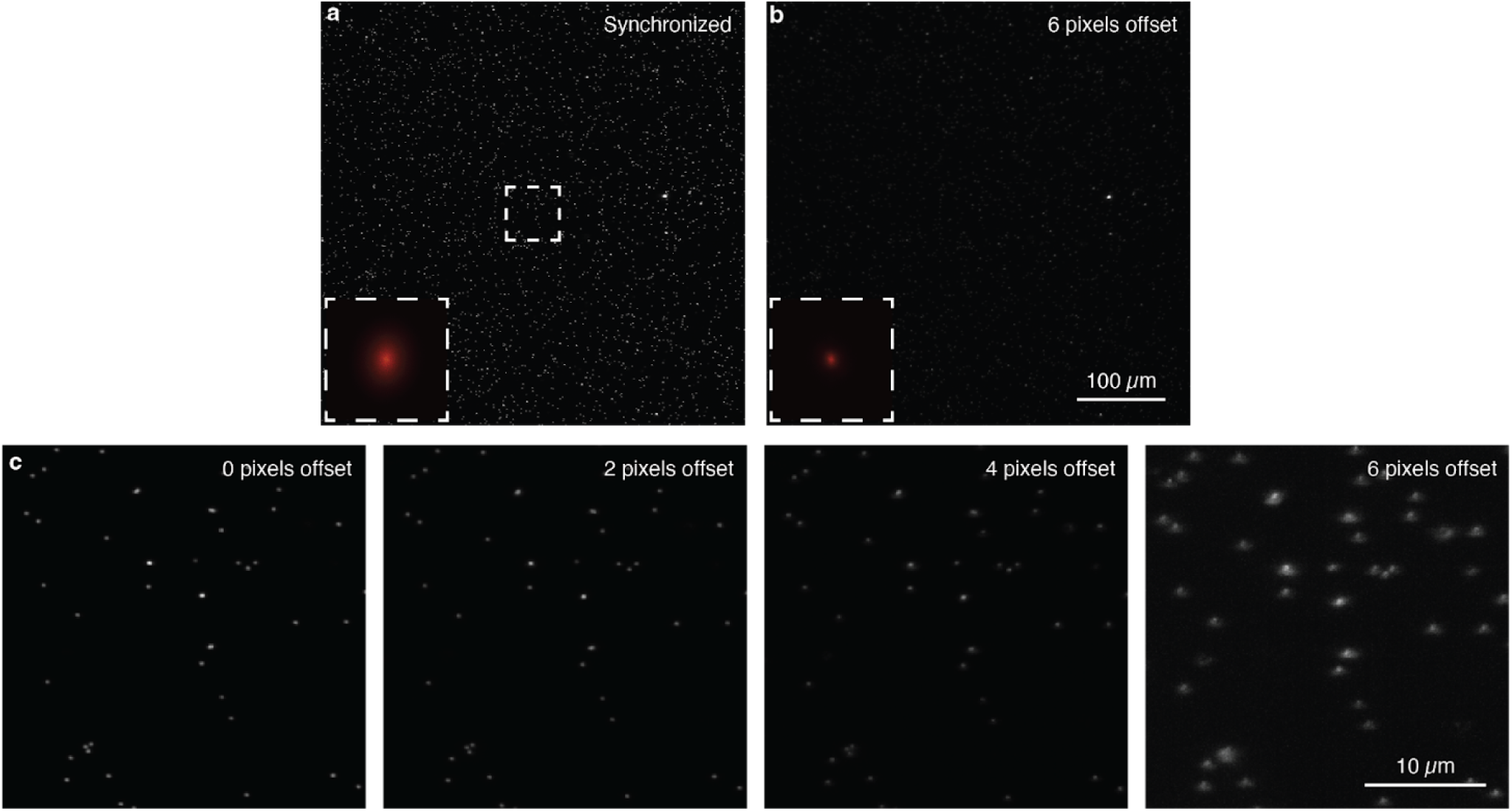
Galvo scanning calibration process. To obtain the best line confocal image quality, the scanning laser line needs to be synchronized to the sCMOS camera’s rolling shutter movement so that the illumination peak falls along the whole scanning line at all times. The calibration process is to determine the voltage to be applied for the galvo mirror at each pixel location to compensate for its non-linear and variable scanning speed of mechanical movement. **a,** The cylindrical lens is rotated by a certain angle θ in the calibration process. **b,** Illustration shows that the laser line projected by the rotated cylindrical lens forms a fixed angle θ in respect to the rolling shutter line on the sCMOS camera sensor during the calibration process. Arrows indicate the scanning directions of the rolling shutter and laser scanning line. The rotated angle blocks most of the laser illumination, resulting only the pixels around their intersection being exposed at any time. When the rolling shutter and the laser line scan across the whole sCMOS chip, their intersection also moves along the scanning direction, resulting in an exposure line along the scanning direction across the whole image frame. **c,** Fluorescent pattern, a straight line perpendicular to the rolling shutter, when there is a perfect synchronization between galvo scanning and the rolling shutter. **d,** Fluorescent pattern when galvo scanning is too rapid. **e,** Fluorescent pattern when galvo scanning is too slow. **f,** Fluorescent pattern when galvo scanning is non-linear. **g-h,** the scanning error before and after the calibration. **i**, Example galvo calibration curves acquired using our automated approach for each channel.

**Fig. S8.**
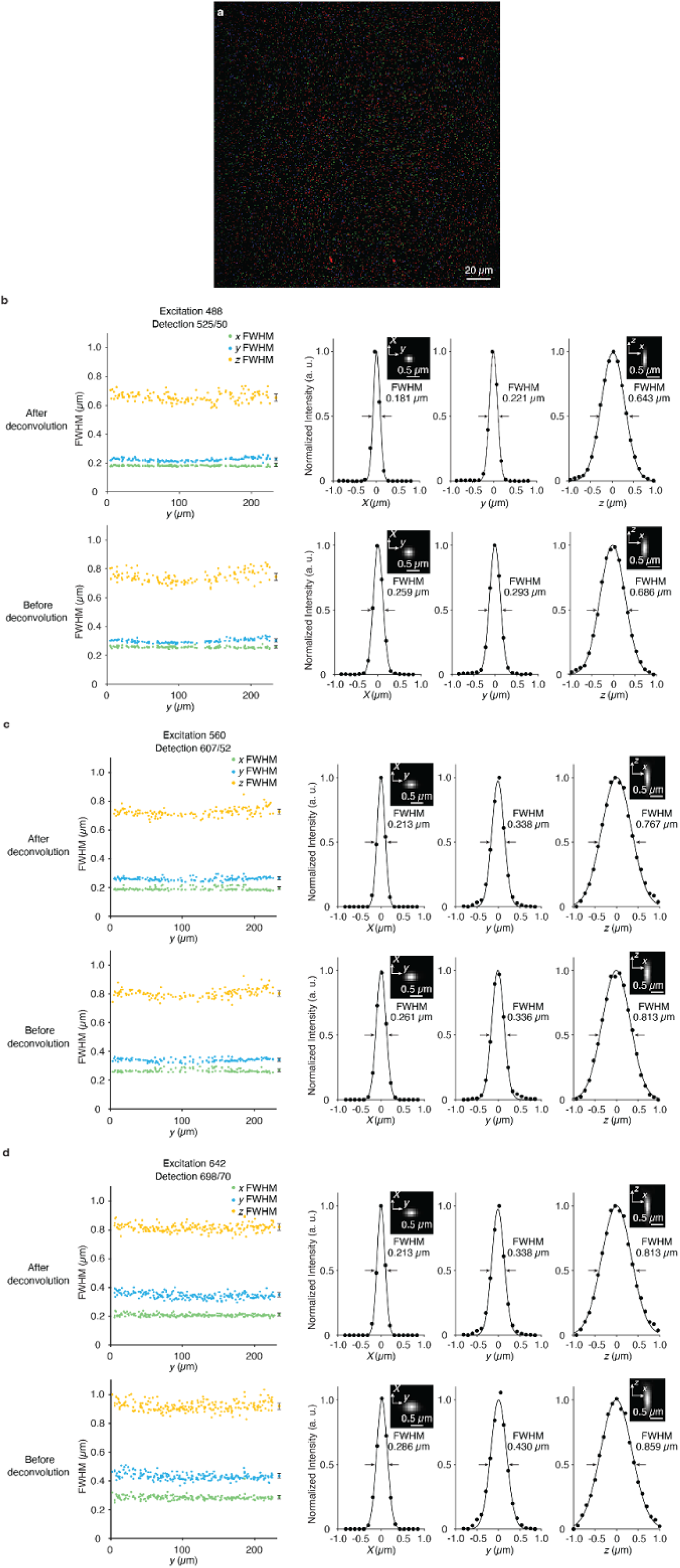
Automated laser excitation drift correction. Due to temperature and humidity change, the galvo position can drift away from the initial calibrated position during a long-duration imaging session, resulting in a parallel shift between the excitation laser line and the rolling shutter. This will cause an intensity drop because part of the excitation laser is blocked by the rolling shutter. We designed a calibration module for laser excitation auto-drift correction (**a**), by which, the power of the excitation laser lights are reduced to a minimum level and the portion transmitting through the master dichroic is directly captured by an auxiliary camera. The excitation laser profile captured right before an experiment will then be used to compute the offset of excitation caused by galvo position drift (**b**). **c,** Captured line patterns on the imaging camera during the correction process. **d,** Enlarged view of the yellow dashed rectangle in **c**.

**Fig. S9.**
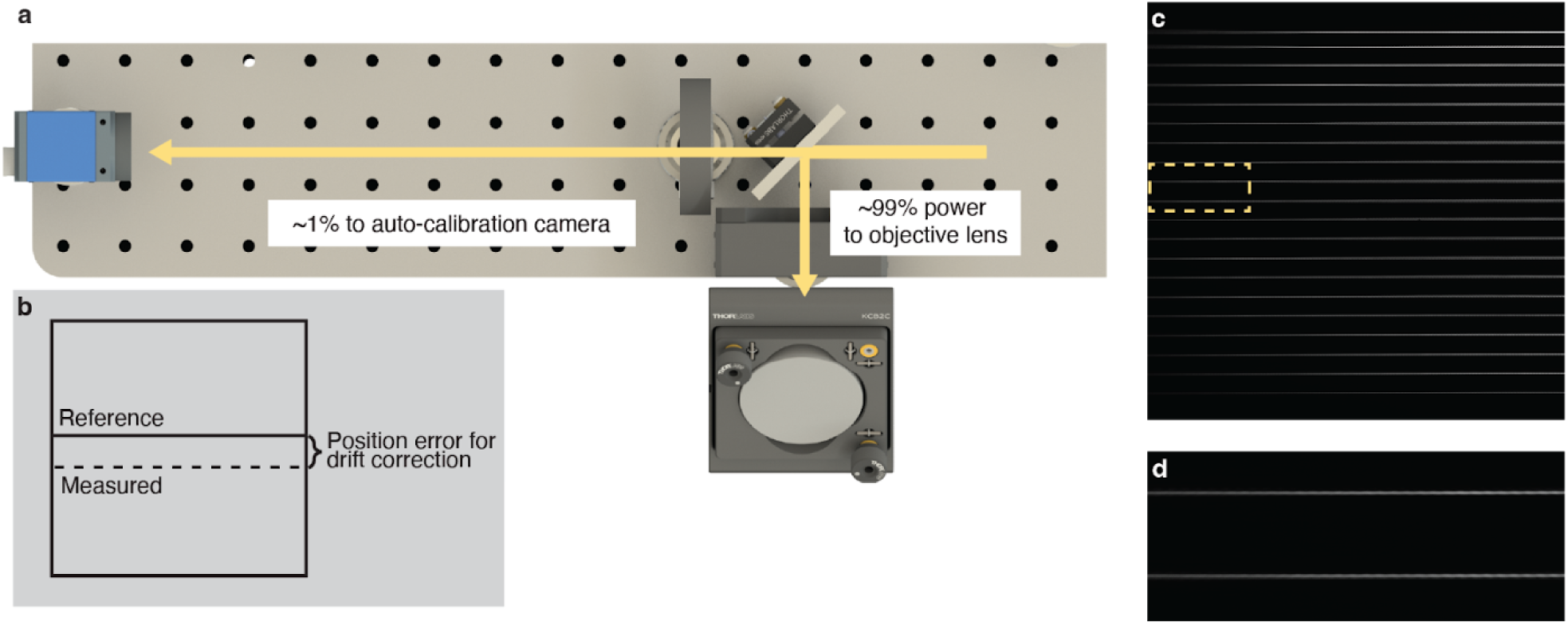
PSF measurement across the imaging volume. a, Maximum-intensity projection (MIP) of 0.2 µm bead images in the 488 nm (**b**), 560 nm (**c**), and 642 nm (**d**) channels. Scale bar, 20 µm. Left columns show the plots of FWHMs, with average FWHMs and standard deviation indicated on the right side of each plot. The plots are presented both before and after 3 rounds of Lucy-Richardson deconvolution with the bead size to correct the influence of bead size on FWHMs. The right columns show the cross-section plot of representative PSF in *x*, *y*, and *z* directions. au, arbitrary units. FWHM, full-width at half-maximum.

**Fig. S10.**
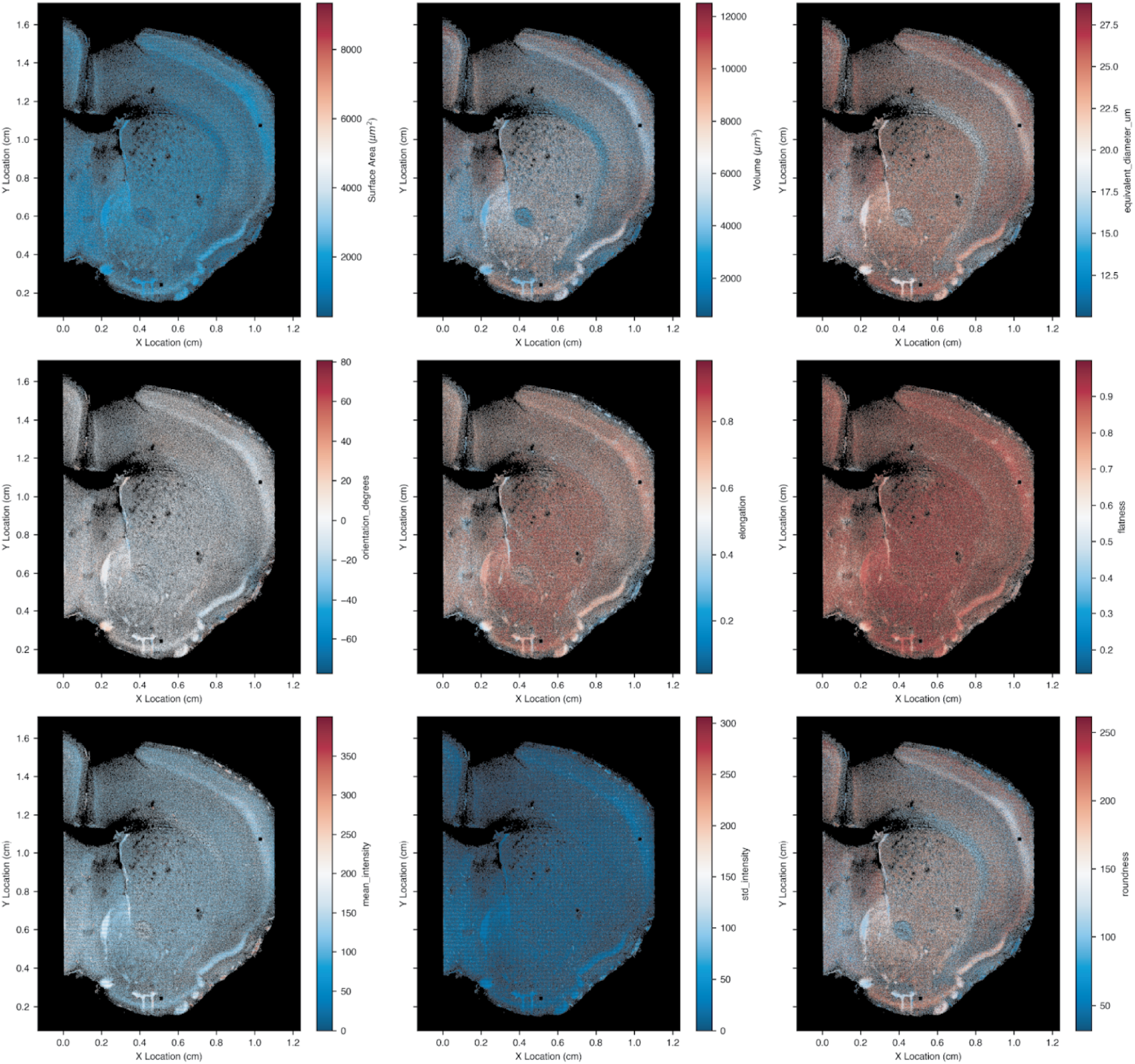
Morphometrics of DRAQ5 nuclei segmentations. Each plot displays a version of Fig. 2g which has been colored by one of the 9 different morphometrics which were calculated from the DRAQ5 stained nuclei segmentation dataset.

**Fig. S11.**
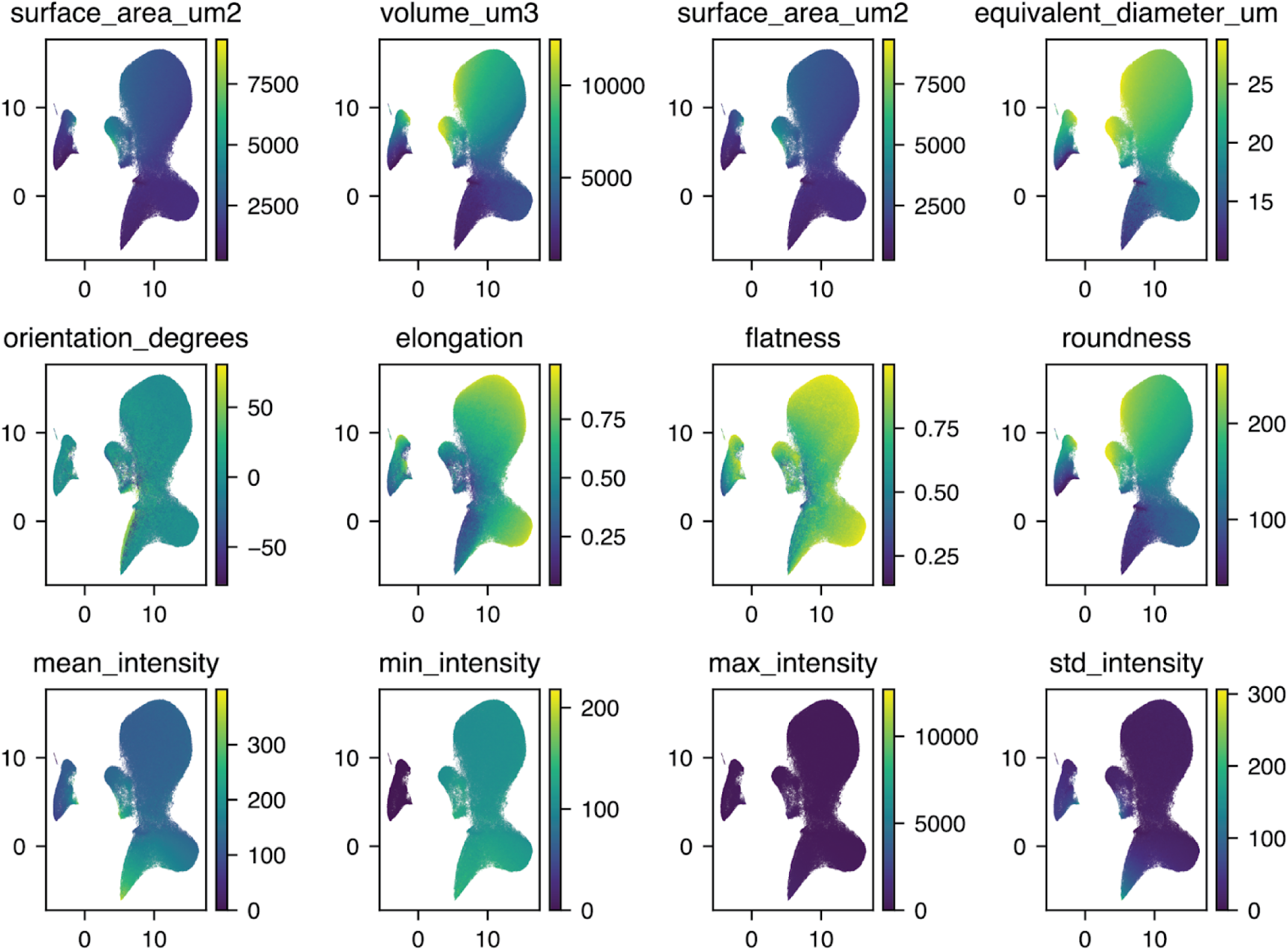
DRAQ5 nuclei segmentation UMAP results. Each plot displays a version of Fig. 2i which has been colored using the same 9 morphometrics displayed in **Fig. S10**, to emphasize the gradients in parameters which are identified in UMAP space.

**Fig. S12.**
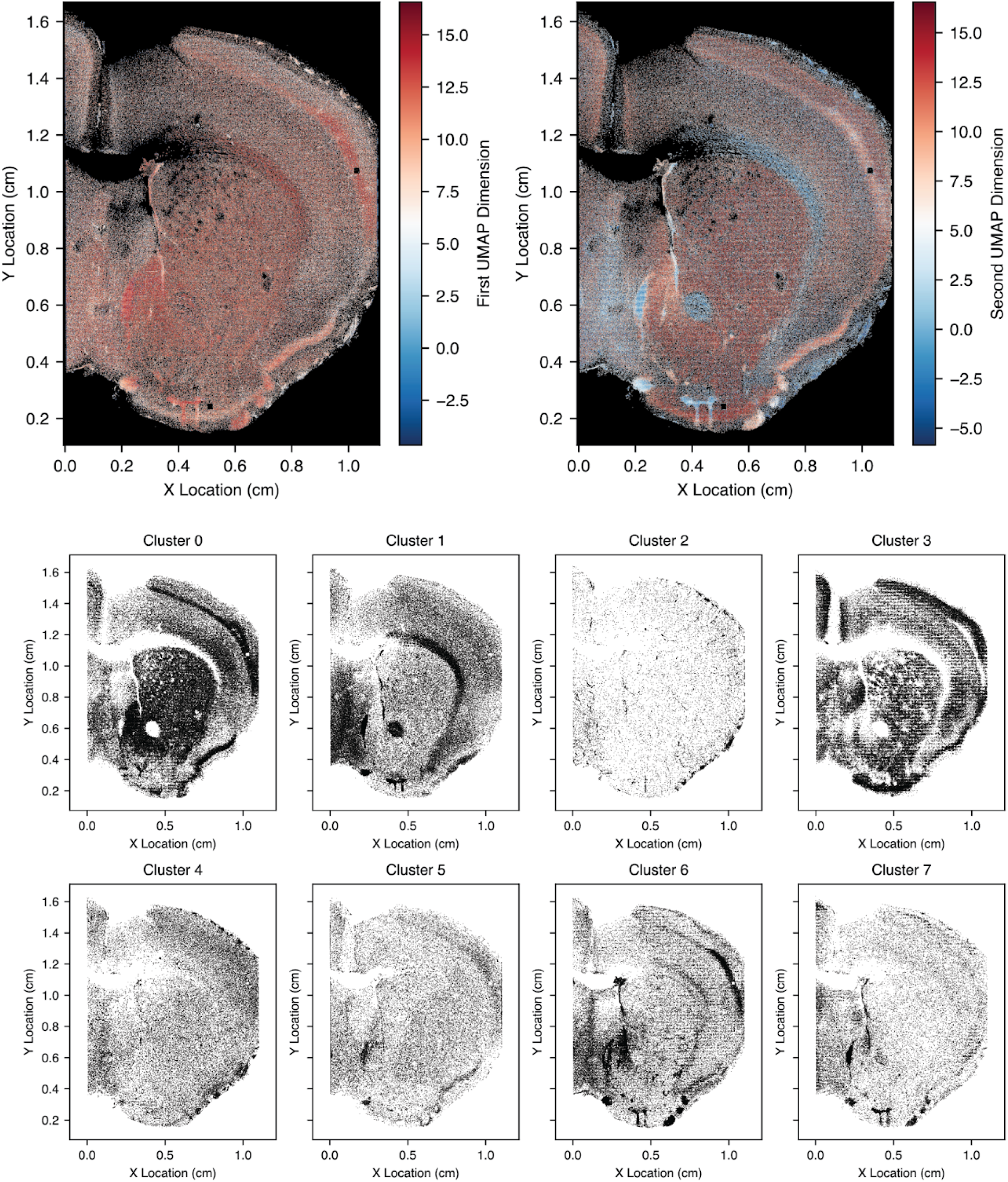
DRAQ5 nuclei segmentation clustering results. Top, scatterplots of the spatial distribution of the UMAP 1st and 2nd dimension. Bottom, scatterplots of the spatial distribution of the 8 GMM cell type clusters identified.

**Fig. S13.**
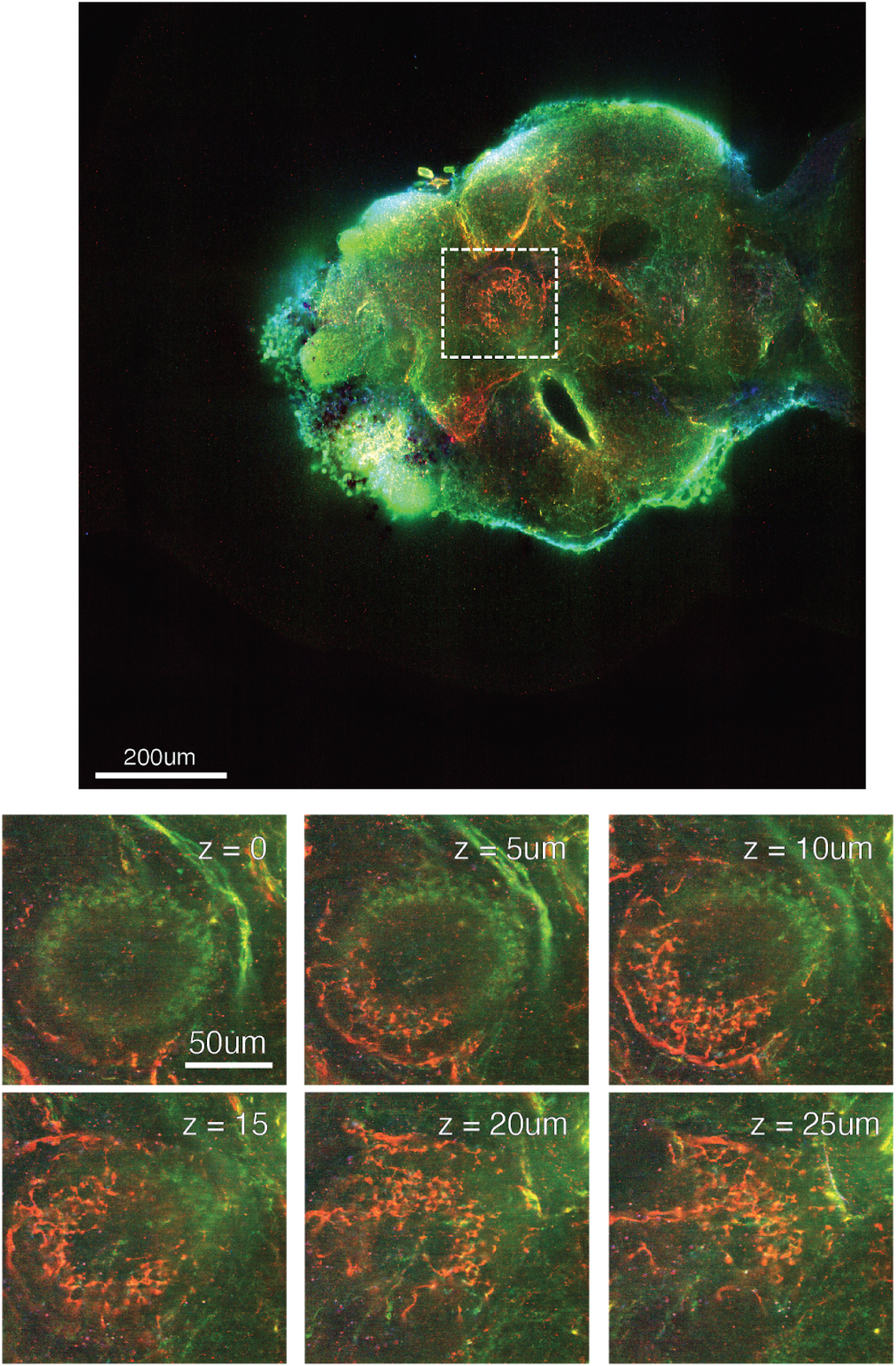
Bitbow *Drosophila* central complex serotonergic neuron. Top, the entire Bitbow fly image with the central complex highlighted in white. Bottom, a Z-frame montage from 0 µm to 25 µm, demonstrating the structure of the central complex innervation by a single serontonergic neuron in Z.

**Fig. S14.**
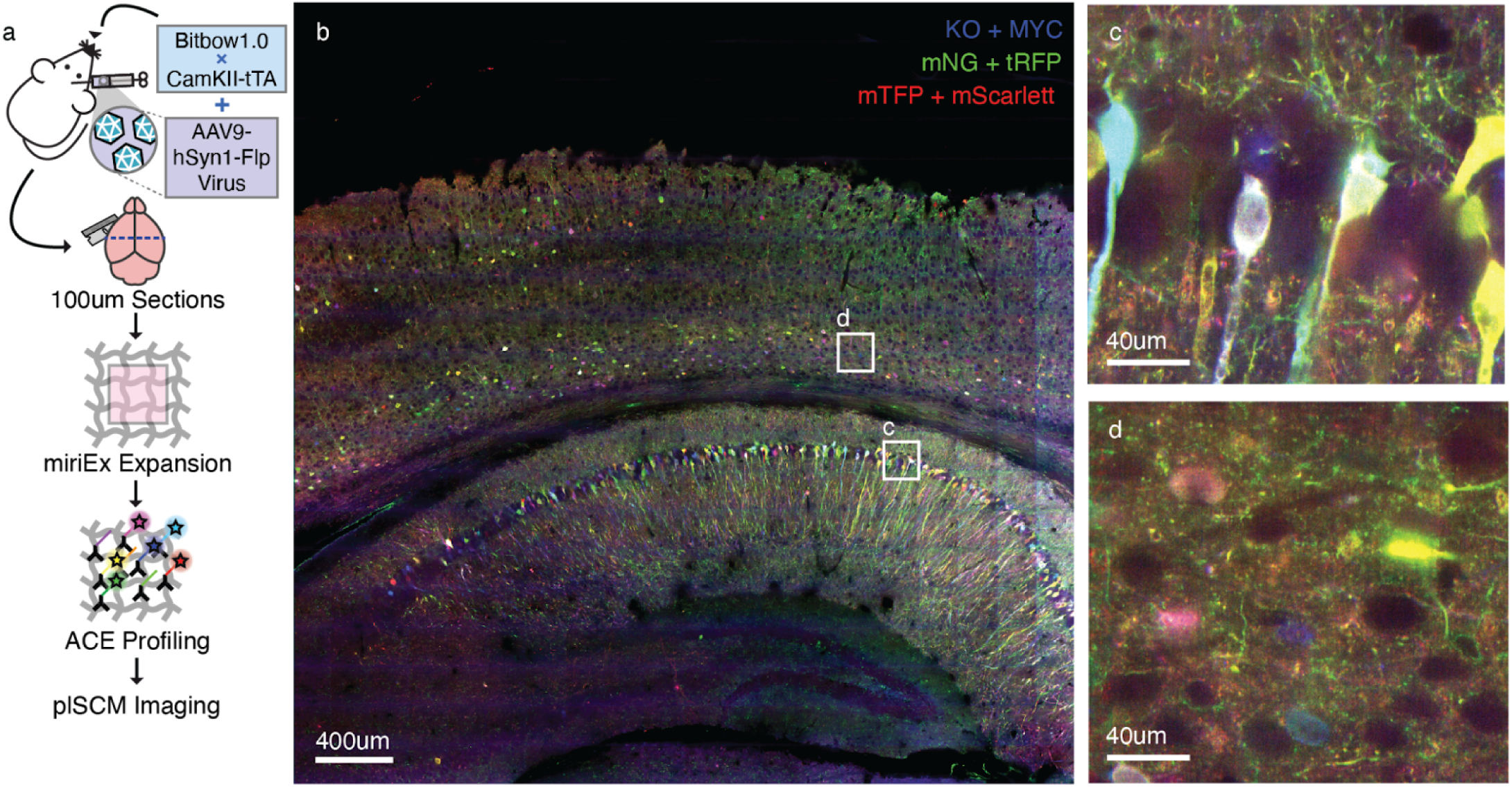
plSCM imaging of Bitbow mouse brain. **a**, Outline of the protocol used to prepare a Bitbow mouse brain sample. In brief, AAV.hSyn-Flp was injected into a Bitbow^+^/CamKII-tTA^+^ young adult mouse. The animal was sacrificed at 5 weeks post injection. Brain section was then gelled and cleared following a miriEx expansion protocol with modification to allow FP secondary antibody amplified by ACE. **b**, Overview of a 26TB image dataset taken with the plSCM. **c,d** Two regions highlighted in **b**, which display multiple colors of Bitbow labeled neurons. AAV, adeno-associated virus; ACE, Amplification by Cyclic Extension.

## Data and Code Availability

Code and design files related to this project is available on GitHub repositories at:

- https://github.com/Cai-Lab-at-University-of-Michigan/plSCMv1_Design
- https://github.com/Cai-Lab-at-University-of-Michigan/plSCMV1_code

## Author Contributions

HJ, LAW, DC, and MC designed and built the plSCM prototype. LAW and DC designed the SNDiF structure, and LAW implemented the SNDiF code. LAW and YL performed microscopy experiments. LAW and BD performed image informatics. YL, XN, MCC, JCH, and KA prepared samples for imaging. HS, KS, and PY developed the ACE labeling strategy and reagents, which was optimized for the miriEx hydrogel by XN and DC. LAW and YL performed neuron tracing experiments, which were further analyzed by BD, RP, AP, SG, and DC. MC and DC conceptualized and managed the project. All authors reviewed and approved this manuscript.

## Acknowledgments

This project was supported by NIH grants RF1MH123402 and RF1MH133764 to DC, and RF1MH124611 to MC and DC. LAW was supported by a University of Michigan Rackham Predoctoral Fellowship. Computation support for this project was provided by University of Michigan Advanced Research Computing (UM-ARC) and the Brain Image Library (BIL) at the Pittsburgh Supercomputing Center (PSC). GPU hardware used was provided by grants from NVIDIA and AMD to LAW and DC. The authors thank Bilyana Popova for the public domain fly drawing used in **Fig. 3**, accessed from bioicons.com.

## Ethics Declarations

A patent US12135411B2 has been granted to HJ and MC describing the plSCM design. LAW and DC are authors of a patent application pub. US20240144418 describing parts of the SNDiF. All animal experiments were performed with the approval of the University of Michigan Institutional Animal Care & Use Committee (IACUC).

## References

1. Walsh, C. L. et al. Imaging intact human organs with local resolution of cellular structures using hierarchical phase-contrast tomography. Nat. Methods 18, 1532–1541 (2021).

2. Susaki, E. A. & Ueda, H. R. Whole-body and whole-organ clearing and imaging techniques with single-cell resolution: Toward organism-level systems biology in mammals. Cell Chem. Biol. 23, 137–157 (2016).

3. Lewis, S. M. et al. Spatial omics and multiplexed imaging to explore cancer biology. Nat. Methods 18, 997–1012 (2021).

4. Kim, G. B. et al. Rapid generation of somatic mouse mosaics with locus-specific, stably integrated transgenic elements. Cell 179, 251–267.e24 (2019).

5. Luo, L. Architectures of neuronal circuits. Science 373, eabg7285 (2021).

6. Muñoz-Castañeda, R. et al. Cellular anatomy of the mouse primary motor cortex. Nature 598, 159–166 (2021).

7. Urai, A. E., Doiron, B., Leifer, A. M. & Churchland, A. K. Large-scale neural recordings call for new insights to link brain and behavior. Nat. Neurosci. 25, 11–19 (2022).

8. Morgan, J. L. & Lichtman, J. W. Why not connectomics? Nat. Methods 10, 494–500 (2013).

9. Eichler, K. et al. The complete connectome of a learning and memory centre in an insect brain. Nature 548, 175–182 (2017).

10. Scheffer, L. K. et al. A connectome and analysis of the adult Drosophila central brain. Elife 9, e57443 (2020).

11. Winding, M. et al. The connectome of an insect brain. Science 379, eadd9330 (2023).

12. Shapson-Coe, A. et al. A connectomic study of a petascale fragment of human cerebral cortex. bioRxiv 2021.05.29.446289 (2021) doi:10.1101/2021.05.29.446289.

13. The MICrONS Consortium, et al. Functional connectomics spanning multiple areas of mouse visual cortex. bioRxiv 2021.07.28.454025 (2023) doi:10.1101/2021.07.28.454025.

14. Winnubst, J. et al. Reconstruction of 1,000 Projection Neurons Reveals New Cell Types and Organization of Long-Range Connectivity in the Mouse Brain. Cell 179, 268–281.e13 (2019).

15. Peng, H. et al. Morphological diversity of single neurons in molecularly defined cell types. Nature 598, 174–181 (2021).

16. Qiu, S. et al. Whole-brain spatial organization of hippocampal single-neuron projectomes. Science 383, eadj9198 (2024).

17. Tavakoli, M. R. et al. Light-microscopy based dense connectomic reconstruction of mammalian brain tissue. bioRxiv 2024.03.01.582884 (2024) doi:10.1101/2024.03.01.582884.

18. Shen, F. Y. et al. Light microscopy based approach for mapping connectivity with molecular specificity. Nat. Commun. 11, 4632 (2020).

19. Cai, D., Cohen, K. B., Luo, T., Lichtman, J. W. & Sanes, J. R. Improved tools for the Brainbow toolbox. Nat. Methods 10, 540–547 (2013).

20. Dean, K. M. et al. Isotropic imaging across spatial scales with axially swept light-sheet microscopy. Nat. Protoc. 17, 2025–2053 (2022).

21. Chhetri, R. K. et al. Whole-animal functional and developmental imaging with isotropic spatial resolution. Nat. Methods 12, 1171–1178 (2015).

22. Voigt, F. F. et al. The mesoSPIM initiative: open-source light-sheet microscopes for imaging cleared tissue. Nat. Methods 16, 1105–1108 (2019).

23. Glaser, A. K. et al. A hybrid open-top light-sheet microscope for versatile multi-scale imaging of cleared tissues. Nat. Methods 19, 613–619 (2022).

24. Yang, B. et al. DaXi-high-resolution, large imaging volume and multi-view single-objective light-sheet microscopy. Nat. Methods 19, 461–469 (2022).

25. Voleti, V. et al. Real-time volumetric microscopy of in vivo dynamics and large-scale samples with SCAPE 2.0. Nat. Methods 16, 1054–1062 (2019).

26. Chen, B.-C. et al. Lattice light-sheet microscopy: imaging molecules to embryos at high spatiotemporal resolution. Science 346, 1257998 (2014).

27. Zimmermann, T. Spectral imaging and linear unmixing in light microscopy. Adv. Biochem. Eng. Biotechnol. 95, 245–265 (2005).

28. Seo, J. et al. PICASSO allows ultra-multiplexed fluorescence imaging of spatially overlapping proteins without reference spectra measurements. Nat. Commun. 13, 2475 (2022).

29. Barentine, A. E. S. et al. An integrated platform for high-throughput nanoscopy. Nat. Biotechnol. 41, 1549–1556 (2023).

30. Zhong, Q. et al. High-definition imaging using line-illumination modulation microscopy. Nat. Methods 18, 309–315 (2021).

31. Walker, L. A., Michki, N. S. & Cai, D. A low-cost and robust microscope hardware trigger interface board. OSF Preprints (2023) doi:10.31219/osf.io/fcb3t.

32. Collet, Y. & Kucherawy, M. S. RFC 8878: Zstandard Compression and the ‘application/zstd’ Media Type. https://www.rfc-editor.org/rfc/rfc8878.

33. Duan, B. et al. Artifact-minimized high-ratio image compression with preserved analysis fidelity. bioRxiv 2024.07. 17.603794 (2024) doi:10.1101/2024.07.17.603794.

34. Truckenbrodt, S. et al. X10 expansion microscopy enables 25-nm resolution on conventional microscopes. EMBO Rep. 19, (2018).

35. Walker, L. A. et al. A browser-based platform for storage, visualization, and analysis of large-scale 3D images in HPC environments. bioRxiv 2024.10.04.616591 (2024) doi:10.1101/2024.10.04.616591.

36. Walker, L. A. et al. nGauge: Integrated and Extensible Neuron Morphology Analysis in Python. Neuroinformatics 20, 755–764 (2022).

37. McInnes, L., Healy, J. & Melville, J. UMAP: Uniform Manifold Approximation and Projection for Dimension Reduction. arXiv [stat.ML] (2018).

38. Livet, J. et al. Transgenic strategies for combinatorial expression of fluorescent proteins in the nervous system. Nature 450, 56–62 (2007).

39. Li, Y. et al. Bitbow Enables Highly Efficient Neuronal Lineage Tracing and Morphology Reconstruction in Single Drosophila Brains. Front. Neural Circuits 15, 732183 (2021).

40. Alekseyenko, O. V., Lee, C. & Kravitz, E. A. Targeted manipulation of serotonergic neurotransmission affects the escalation of aggression in adult male Drosophila melanogaster. PLoS One 5, e10806 (2010).

41. Dorkenwald, S. et al. FlyWire: online community for whole-brain connectomics. Nat. Methods 19, 119–128 (2022).

42. Peng, T. et al. A BaSiC tool for background and shading correction of optical microscopy images. Nat. Commun. 8, 14836 (2017).

43. Tange, O. GNU parallel: The command-line power tool. login - The Usenix Magazine 36, (2011).

44. Hörl, D. et al. BigStitcher: reconstructing high-resolution image datasets of cleared and expanded samples. Nat. Methods 16, 870–874 (2019).

45. Pizer, S. M., Eugene Johnston, R., Ericksen, J. P., Yankaskas, B. C. & Keith E. Muller Medical Image Display Research Group. Contrast-Limited Adaptive Histogram Equalization: Speed and Effectiveness. Preprint at http://www.cs.unc.edu/techreports/90-035.pdf.

46. Isensee, F., Jaeger, P. F., Kohl, S. A. A., Petersen, J. & Maier-Hein, K. H. nnU-Net: a self-configuring method for deep learning-based biomedical image segmentation. Nat. Methods 18, 203–211 (2021).

47. Isensee, F., et al. nnInteractive: Redefining 3D Promptable Segmentation. arXiv [cs.CV] (2025).

48. Pedregosa, F. et al. Scikit-learn: Machine Learning in Python. J. Mach. Learn. Res. 12, 2825–2830 (2011).

49. Bogovic, J. A. et al. An unbiased template of the Drosophila brain and ventral nerve cord. PLoS One 15, e0236495 (2020).

50. Court, R. et al. Virtual Fly Brain-An interactive atlas of the Drosophila nervous system. Front. Physiol. 14, 1076533 (2023).

51. Schlegel, P. et al. Whole-brain annotation and multi-connectome cell typing of Drosophila. Nature 634, 139–152 (2024).

52. Takeshita, N. et al. Acto3D: an open-source user-friendly volume rendering software for high-resolution 3D fluorescence imaging in biology. Development 151, (2024).

53. Schindelin, J., et al. Fiji: an open-source platform for biological-image analysis. Nat. Methods 9, 676–682 (2012).

