## Supplementary material for "A parallelly distributed microscope and software system for scalable high-throughput multispectral 3D imaging": Supp Video Legends

**Supplementary Videos**

**Supp. Video 1 | A z-axis flythrough of GFP-labeled pyramidal neurons.** A cortical region of the image in **Fig. 2b** was rendered using Neuroglancer, showing the GFP native fluorescence in a 2.7-fold expanded GFP-M transgenic mouse brain section.

**Supp. Video 2 | 3D volumetric rendering of GFP-labeled pyramidal neurons, GFAP-stained astrocytes, and DRAQ5-stained nuclei.** The cortical region highlighted in **Fig. 2c** was rendered using Acto3D. Yellow represents the DRAQ5+ nuclei, cyan represents GFP+ neurons, and magenta represents the GFAP+ astrocytes. Contrast and gamma adjustments were performed to improve the visibility of the rendering.

**Supp. Video 3 | 3D volumetric rendering of Bitbow-labeled 4-fold expanded *Drosophila* brain.** A downsampled version of the data displayed in **Fig. 3c** was rendered using Acto3D. The three color channels are mapped to the corresponding color channels identified in **Fig. 3c**. Contrast and gamma adjustments were applied to enhance the visibility of the rendering.

**Supp. Video 4 | 3D volumetric rendering of Bitbow-labeled 3-fold expanded mouse section.** A downsampled version of the data in **Fig. S14b** was rendered using Acto3D. Maximum projection rendering is used to display as many neurons as possible without occlusion. Contrast and gamma adjustments were performed to improve the visibility of the rendering.

**Supp. Video 5 | A z-axis flythrough of LICONN-processed ~10-fold expanded mouse cortical section.** A downsampled version of the data displayed in **Fig. 4b** was rendered using the Fiji video export mode, followed by contrast adjustment and conversion with VLC.

**Supp. Video 6 | A z-axis flythrough of zoomed-in, LICONN-processed ~10-fold expanded mouse cortical section.** A full-resolution subset of the image in **Fig. 4b** was downloaded and rendered using the Fiji video export mode, followed by contrast adjustment and conversion with VLC.
